# COA8 is a COX10-binding protein involved in the early biogenesis of cytochrome *c* oxidase

**DOI:** 10.1101/2024.04.02.587738

**Authors:** Michele Brischigliaro, Kristýna Čunátová, Alfredo Cabrera-Orefice, Jimin Pei, Cinzia Franchin, Marco Roverso, Suleva Povea-Cabello, Sara Bogialli, Giorgio Arrigoni, Susanne Arnold, Qian Cong, Massimo Zeviani, Carlo Viscomi, Erika Fernández-Vizarra

**Affiliations:** Department of Biomedical Sciences, University of Padova, Padova, Italy; Veneto Institute of Molecular Medicine, Padova, Italy; Radboud Institute for Molecular Life Sciences, Radboud University Medical Center, Nijmegen, The Netherlands; Functional Proteomics Center, Institute for Cardiovascular Physiology, Goethe University Frankfurt, Frankfurt am Main, Germany; Eugene McDermott Center for Human Growth and Development, Department of Biophysics, and Harold C. Simmons Comprehensive Cancer Center, University of Texas Southwestern Medical Center, Dallas, TX, USA; Department of Chemical Sciences, University of Padova, Padova, Italy; Cologne Excellence Cluster on Cellular Stress Responses in Aging-Associated Diseases (CECAD), University of Cologne, Cologne, Germany; IRCCS Women and Children’s Hospital “Burlo-Garofolo”, Trieste, Italy; Department of Neurosciences, University of Padova, Padova, Italy

**Keywords:** mitochondrial respiratory chain, complex IV assembly, cytochrome c oxidase, assembly factor, COA8, COX10, heme A synthesis, heme O synthase

## Abstract

Human cytochrome *c* oxidase (COX) is composed of fourteen subunits, only two of which are catalytic. The functions of a myriad of accessory factors are needed to put together the structural subunits, as well as for the formation of the active centers. The molecular roles of most of these factors are still unknown, even if they are essential for COX function since their defects cause severe mitochondrial disease. *COA8* is exclusively found in metazoa and loss-of-function genetic variants cause mitochondrial encephalopathy and COX deficiency in humans. In this work, we have resolved the function of COA8, as a factor that specifically binds to COX10, the heme O synthase. This interaction is necessary not only for the stabilization of human COX10, but also to guarantee its catalytic activity as the first step for the synthesis of the heme A cofactor, which is a crucial component of the COX catalytic center.

## INTRODUCTION

Mitochondrial diseases are a group of highly heterogeneous syndromes, mainly characterized by impairment of oxidative phosphorylation (OXPHOS) (Gorman *et al*, 2016). The mammalian OXPHOS system is composed of the four respiratory chain complexes (I-IV), two mobile electron carriers (coenzyme Q and cytochrome c) and the ATP synthase (complex V). The respiratory chain, or electron transfer system (ETS), carries out a series of redox reactions, transporting electrons throughout the inner mitochondrial membrane (IMM), to finally reduce molecular oxygen to water, a reaction catalyzed by the cytochrome *c* oxidase (complex IV). Electron transfer is coupled to proton pumping from the matrix to the intermembrane space across the IMM. Proton translocation is operated by complexes I, III, and IV, and creates a respiration-induced electrochemical gradient. The proton-motive force produced by mitochondrial respiration, sustains the condensation of ADP and inorganic phosphate into ATP through the proton-driven rotational motor of the ATP synthase, completing the bioenergetic OXPHOS process. This is a very efficient process, which converts energy stored in nutrients into the common energy currency of the cells, i.e. ATP (Vercellino & Sazanov, 2022). Either of the respiratory complexes can be defective and cause mitochondrial dysfunction. To date, pathological variants in more than 400 genes have been described as causative of primary mitochondrial disease (Schlieben & Prokisch, 2023). Genes encoding either structural OXPHOS components or assembly factors, necessary for the biogenesis of each complex but not part of their final structures, constitute a large portion of these disease-associated genes (Fernandez-Vizarra & Zeviani, 2021b). Cytochrome *c* Oxidase Assembly factor 8 (COA8) is a protein that assists the biogenesis of complex IV (cIV), the function of which was unknown before the discovery that loss-of-function mutations in its gene cause mitochondrial disease with cytochrome *c* oxidase (COX) deficiency in both humans and animal models (Brischigliaro *et al*, 2023; Brischigliaro *et al*, 2019; Melchionda *et al*, 2014; Signes *et al*, 2019). Pathological variants in *COA8* (previously known as *APOPT1*) have been described in nine individuals from eight different families, associated with cavitating leukoencephalopathy and mitochondrial cIV deficiency (OMIM #619061) (Chapleau *et al*, 2023; Hedberg-Oldfors *et al*, 2020; Melchionda *et al*., 2014; Sharma *et al*, 2018) and more recently in one family presenting with myopathy as the main clinical feature (Rimoldi *et al*, 2023). CIV (or COX) is a heme-copper oxygen reductase that performs the terminal redox reaction of cellular respiration, i.e., oxidation of cytochrome *c* and reduction of molecular oxygen to water (Sousa *et al*, 2012; Wikstrom *et al*, 2018). Human cIV is composed of three mitochondrial DNA-encoded core subunits and eleven nuclear-encoded supernumerary subunits (Cunatova *et al*, 2020). The assembly of human cIV subunits is hierarchically organized (Nijtmans *et al*, 1998), and recent evidence indicates that it proceeds in a modular fashion, with each of the modules defined by each one of the three mtDNA-encoded core subunits, MT-CO1, MT-CO2 and MT-CO3 (Vidoni *et al*, 2017). Complex IV can be found either as an individual entity (cIV monomer), as a dimer (cIV_2_) or associated with dimeric complex III (cIII_2_) in different supercomplex (SC) types, either alone (SCIII_2_+IV) or bound to complex I, forming the SCI+III_2_+IV, also known as the respirasome (Schagger & Pfeiffer, 2000; Vercellino & Sazanov, 2022). Evidence indicates that the pathways for the full assembly of the individual cIV or SC-associated cIV might be different (Fernandez-Vizarra *et al*, 2022; Fernandez-Vizarra & Ugalde, 2022; Lobo-Jarne *et al*, 2020; Timon-Gomez *et al*, 2020). The synthesis and initial assembly of MT-CO1 is a particularly intricate and highly regulated process, in which a plethora of additional assembly factors are involved (Dennerlein *et al*, 2023). However, a large number of proteins are also necessary to assist the maturation and incorporation of the remaining subunits required to build up a functional cIV (Timon-Gomez *et al*, 2018). An important group of cIV assembly factors are involved in the synthesis and insertion of the metal cofactors into the catalytic centers of the MT-CO1 and MT-CO2 apoproteins (Nyvltova *et al*, 2022). MT-CO2 contains a Cu_A_ diatomic center, whereas MT-CO1 contains a Cu_B_ atom and two heme A moieties (cytochrome *a* and *a3*), forming the binuclear center where oxygen reduction occurs (Wikstrom *et al*., 2018). Heme A synthesis is a two-step reaction that occurs within mitochondria: heme B is first farnesylated by the heme O synthase denominated COX10. Subsequently, COX15, or heme A synthase, catalyzes the conversion of heme O into the final product, heme A (Swenson *et al*, 2020). Mutations in either gene encoding COX10 or COX15 have been associated with severe mitochondrial disease related with cIV deficiency (Antonicka *et al*, 2003a; Antonicka *et al*, 2003b; Diaz *et al*, 2005; Viscomi *et al*, 2011). The interaction of COX10 with COA8 was predicted by computational analysis (Pei *et al*, 2022), but the experimental demonstration of this interaction and its pathophysiological relevance has been addressed in this work.

Here, we show that COA8 plays an essential role at the earliest steps of cIV biogenesis by being involved in heme A synthesis through the specific interaction with COX10, which is crucial for MT-CO1 functional maturation. In addition, we have established that binding of COA8 is not only necessary for the structural stabilization of COX10, but also has an additional functional role in which the COA8 highly conserved W174, located in the interaction surface between the two proteins, is necessary for maintaining physiological heme A levels and cIV assembly and activity.

## RESULTS

### Absence of COA8 impairs early stages of cIV assembly

The biochemical characterization of patient samples established that the mitochondrial respiratory chain defect associated with *COA8* loss-of-function mutations is an isolated cytochrome *c* oxidase (cIV) defect (Hedberg-Oldfors *et al*., 2020; Melchionda *et al*., 2014). The analysis of the cIV assembly state in patient-derived fibroblasts and in tissues of mouse and fruit fly Coa8 knock-out (KO) models, showed significantly lower amounts of fully assembled cIV and the accumulation of partially assembled intermediates (Brischigliaro *et al*., 2023; Signes *et al*., 2019). In order to pinpoint the precise step in which cIV assembly is stalled when COA8 is absent, we used the same strategy that allowed us to characterize the cIV assembly intermediates accumulated in a *MT-CO3* defective cell line (Vidoni *et al*., 2017). Thus, we measured the relative amounts of immunocaptured cIV subunits in patient-derived immortalized fibroblasts, obtained from individual S6 (Melchionda *et al*., 2014) (expressing GFP as a negative control—S6+GFP), and compared it with that obtained in the same cell line expressing GFP-tagged wild-type (WT) COA8 sequence (S6+COA8^GFP^), completely reverting the cIV defect (Signes *et al*., 2019). The quantitative proteomics analyses were based on isotope labeling of amino acids in cell culture (SILAC) and cIV immunocapture from mitoplasts extracted from a 1:1 mixture of differentially labelled S6+GFP and S6+COA8^GFP^ cells (**Figure 1A**). The experiment was performed in duplicate with reciprocally labeled samples and the results of each replicate (**Supplemental Table S1**) were plotted in the x– and y-axis, respectively, in the graph shown in **Figure 1B**. These results indicated that the levels of the COX4I1 and COX5A subunits and of the cIV-associated protein HIGD1A, were unchanged between the pathological (S6+GFP) and functional (S6+COA8^GFP^) cells. In contrast, the levels of the remaining cIV structural subunits were decreased to the same extent in the COA8-null cells compared with the control (S6+COA8^GFP^). Therefore, the defect produced by the absence of the COA8 factor occurs at early steps of the cIV assembly pathway (**Figure 1C**), most likely at the level of maturation and/or incorporation of MT-CO1, since the synthesis of this mtDNA-encoded apoprotein was not affected (Signes *et al*., 2019).

**Figure 1.**
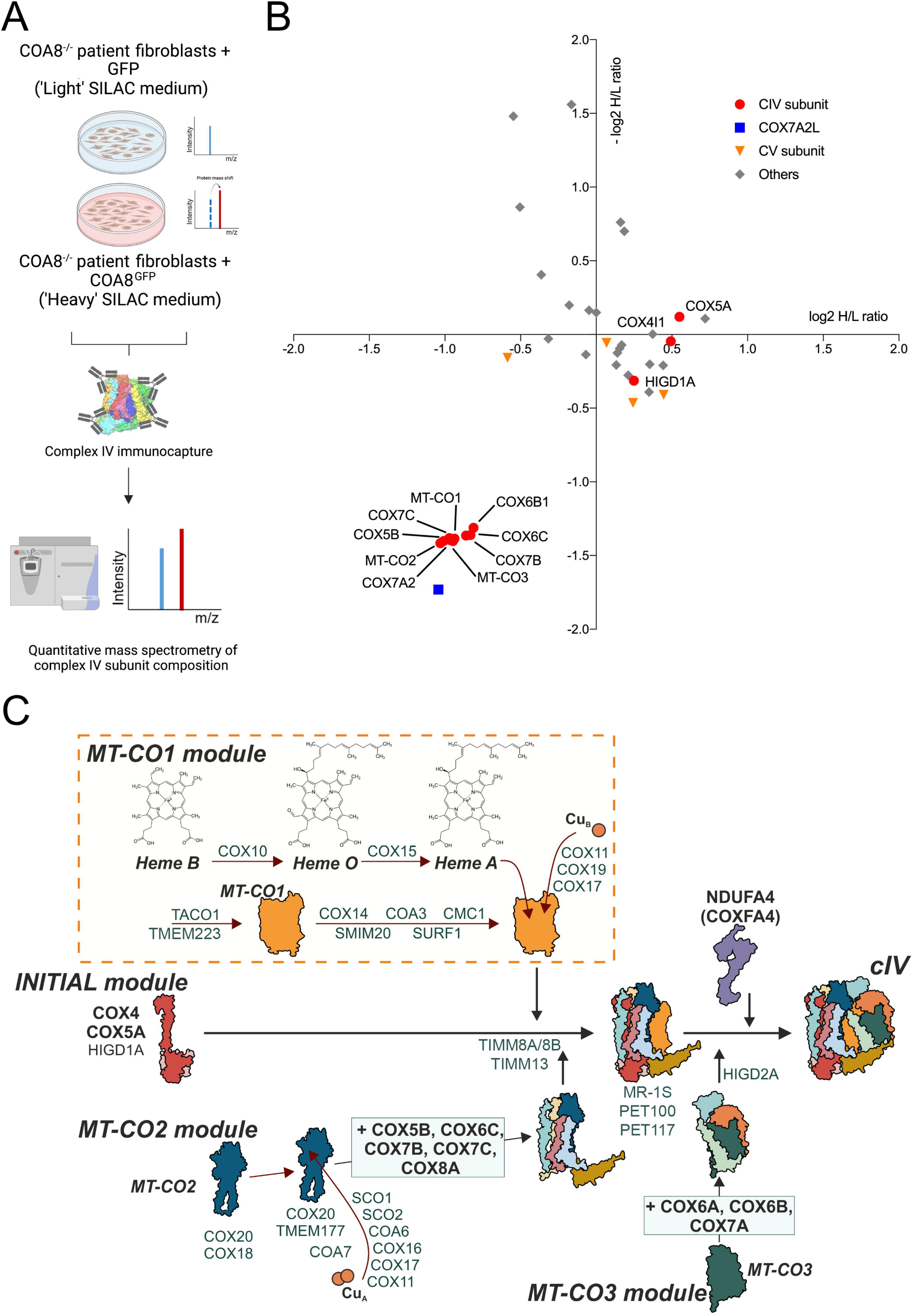
The absence of COA8 impairs early stages of cIV assembly. (**A**) Workflow of SILAC-based quantitative proteomic analysis of complex IV in uncomplemented (S6+GFP) and complemented (S6+COA8^GFP^) patient fibroblasts. Cells are grown either in ‘light’ medium, containing non-labeled amino acids, or in ‘heavy’ medium, containing amino acids labeled with non-radioactive isotopes (^13^C and ^15^N). Cells were mixed at 1:1 ratio, and mitochondria-enriched fractions were subjected to complex IV immunocapture and analyzed by quantitative MS to identify subunit composition of fully assembled complex IV and sub-assembly intermediates. **(B)** Reverse SILAC-labeled mitochondria-enriched fractions from immortalized fibroblasts were isolated, mixed, and subjected to complex IV immunocapture. The values in the x axis correspond to the log2 heavy-to-light (H/L) ratio of the proteins detected in experiment 1, where the heavy (H)-labeled complemented (S6+COA8^GFP^) patient fibroblasts and unlabeled (L) uncomplemented (GFP) patients’ fibroblasts were mixed. The values in the y axis correspond to the inverted log2 H/L ratio (−log2 H/L) of the proteins detected in experiment 2, where the unlabeled (L) complemented (COA8-GFP) patient fibroblasts and labeled (H) uncomplemented (GFP) patients’ fibroblasts were mixed. Red dots = complex IV subunits; blue square = COX7A2L subunit (aka SCAFI); orange triangles = complex V subunits; gray squares = others. **(C)** Schematic representation of the current model for the ‘monomeric’ complex IV assembly pathway, based on (Vidoni *et al*., 2017), indicating the relevant assembly factors involved in each step. Cartoons derive from published human cryo-EM structures deposited with PDB ID 5Z62.

### COA8 specifically interacts with the heme O synthase COX10

Next, we sought to better understand the molecular function of COA8 by determining its interactome. We took advantage of the SILAC approach allowing us to discern contaminants and unspecific interactions from the specific COA8 interactors amongst the proteins identified by mass spectrometry (MS) in the co-immunopurified fractions. For this experiment, we used 143B cells stably expressing an HA-tagged version of COA8 (Signes *et al*., 2019), whereas the negative controls were 143B cells transduced with GFP. The MS analysis of the co-immunopurified fractions from the duplicate experiments (inverting the labeling), revealed a very specific interaction of COA8 with COX10, being these the only two proteins significantly enriched (**Figure 2A**, **Supplemental Table S2**). COA8 is a ROS-responsive protein, since its intra-mitochondrial levels are increased in response to oxidative stress (Signes *et al*., 2019). However, the exclusive interaction with COX10 was maintained in the presence of H_2_O_2_ (**Supplemental Figure S1**, **Supplemental Table S2**). Additionally, the COA8-COX10 interaction was confirmed by co-immunopurification of COA8^GFP^ from 143B cells using an anti-GFP nanobody (**Figure 2B**).

**Figure 2.**
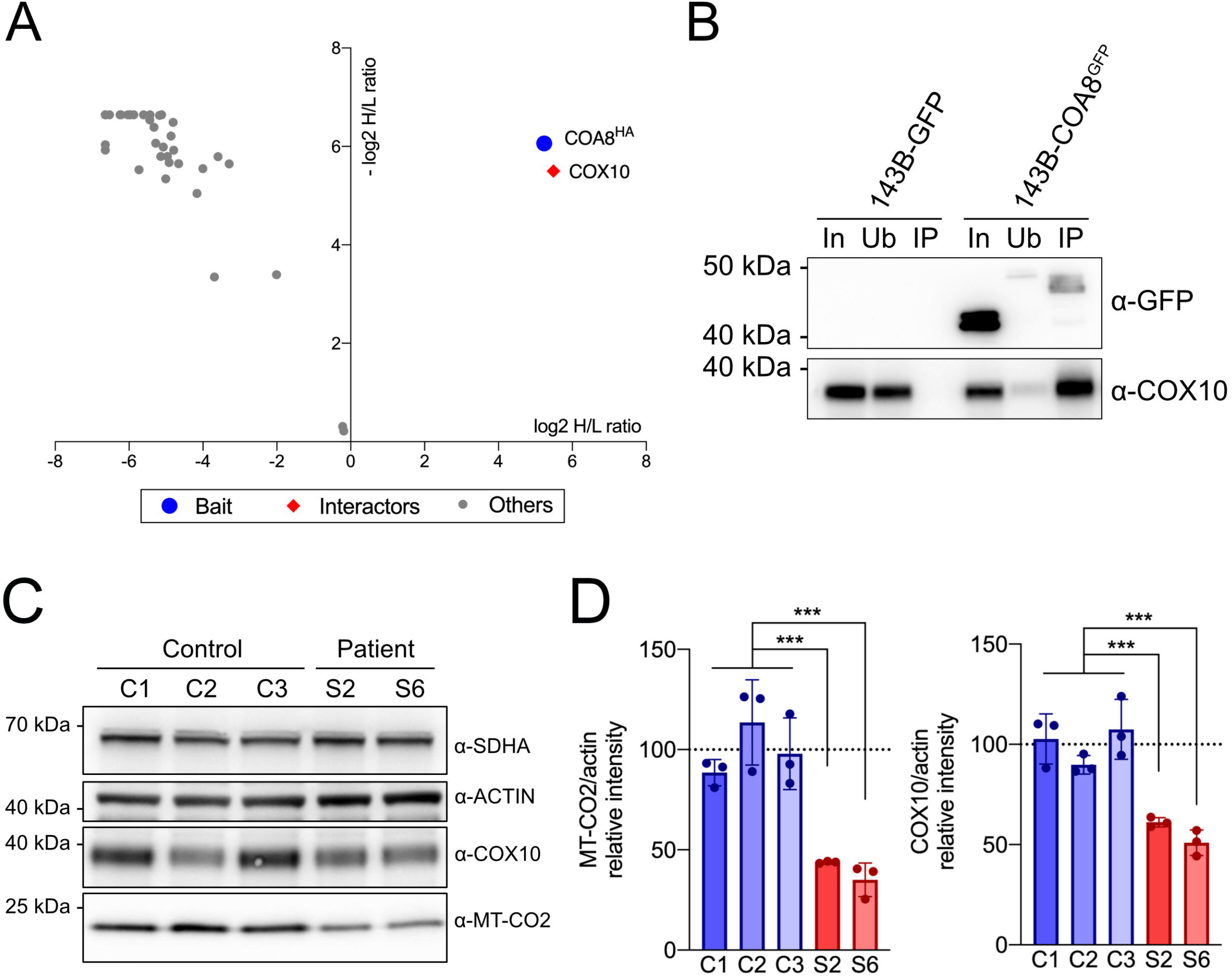
COA8 specifically interacts with the heme O synthase COX10 and stabilizes it. (**A**) Reverse SILAC-labeled mitochondria-enriched fractions from 143B osteosarcoma cells either expressing HA-tagged COA8 (COA8^HA^) or GFP were isolated, mixed, and subjected to anti-HA immunopurification. The values in the x axis correspond to the log2 heavy-to-light (H/L) ratio of the proteins detected in experiment 1, where the heavy (H)-labeled COA8^HA^ expressing cells and unlabeled (L) GFP expressing cells were mixed. The values in the y axis correspond to the inverted log2 H/L ratio (−log2 H/L) of the proteins detected in experiment 2, where the unlabeled (L) COA8^HA^ expressing cells and labeled (H) GFP expressing cells were mixed. Blue dot = bait (HA); red squares = interactors; grey dots = others. **(B)** Western blot analysis of anti-GFP affinity purified complexes from 143B cells expressing either GFP or COA8^GFP^. In = total mitochondrial-enriched fractions (input); Ub = unbound material; IP = immunoprecipitated material. Samples were probed with antibodies against GFP and COX10. **(C)** Steady state levels of MT-CO2, COX10, Actin and SDHA in three control immortalized skin fibroblast cell lines (C1-3) and in patient-derived immortalized skin fibroblasts from two unrelated subjects carrying loss-of-function variants in COA8 (S2, S6). **(D)** Densitometric analysis of MT-CO2 and COX10 levels normalized by ACTIN in healthy control cell lines (C1-3) and 2 unrelated COA8 subjects (S2, S6). Data are plotted as mean ± S.D. (n = 3 biological replicates, one-way ANOVA with Tukey’s multiple comparisons, **p ≤ 0.01, ***p ≤ 0.001).

These results prompted us to check the steady-state levels of COX10 in two immortalized fibroblast cell lines derived from two unrelated patients carrying loss-of-function variants in *COA8* (subjects S2 and S6 (Melchionda *et al*., 2014; Signes *et al*., 2019)). As shown in **Figure 2C** and **Figure 2D**, the absence of COA8 was associated with the decrease in the cIV subunit MT-CO2, and also with significantly lower abundance of COX10. This is in agreement with the observation that COA8 and COX10 specifically interact with each other, likely stabilizing COX10 structure/function.

### COA8 is predicted to interact with the COX10 central cavity catalytic domain through a set of conserved residues

Based on large-scale coevolution analysis and deep learning-based structure modeling, the interaction between COA8 and COX10 has been predicted with a high degree of confidence (Pei *et al*., 2022). COX10 (443 amino acids) presumably contains nine transmembrane helices, whereas COA8, a smaller protein (206 amino acids), is predicted to fold into two anti-parallel alpha helices (**Figure 3A**). The contact between COA8 and COX10 was predicted to occur mainly between the second alpha helix of COA8 and a helix loop (HL) located between the second and third transmembrane domains (HL2-3) of COX10 (Pei *et al*., 2022). This HL2-3 spans from position 195 to position 234 in the COX10 amino acid sequence (**Figure 3B**) and forms a central pocket, containing essential residues for the heme O synthase activity (Rivett *et al*, 2021), which are highly conserved from bacteria to human (**Figure 3B**). Predicted contacts with score above 0.9 (**Table 1**, **Supplemental Table S3**) were considered to predominantly contribute to the interaction between the two proteins. These contacts involve several large hydrophobic amino acids in COA8, which are mainly aromatic residues. Based on the probability of contact and on the conservation among metazoan species (**Figure 3C**), the only kingdom in which COA8 homologs exist, we determined that several of these hydrophobic/aromatic residues were likely relevant for the interaction with COX10, namely: W111, F118, F159, L160, Y170, W174 and Y175. These residues are located in the COA8 domain were confidently predicted to interact with COX10 residues mostly present in the catalytic pocket domain (**Figure 3A**, **Figure 3B** and **Table 1, Supplemental Table S3**). Interestingly, the proline residue in position 213 (P213) in human COX10 has a very high contact prediction score with W174 of COA8 (**Table 1** and **Supplemental Table S3**) and among all the residues considered to be fundamental for catalysis in COX10, present in the HL2-3 (Rivett *et al*., 2021), P213 is the only one that is not conserved from bacteria to humans. In fact, this position is occupied by a Pro residue only in metazoans, whereas in bacteria and yeast it is an acidic residue (either D or E) (**Figure 3B**).

**Figure 3.**
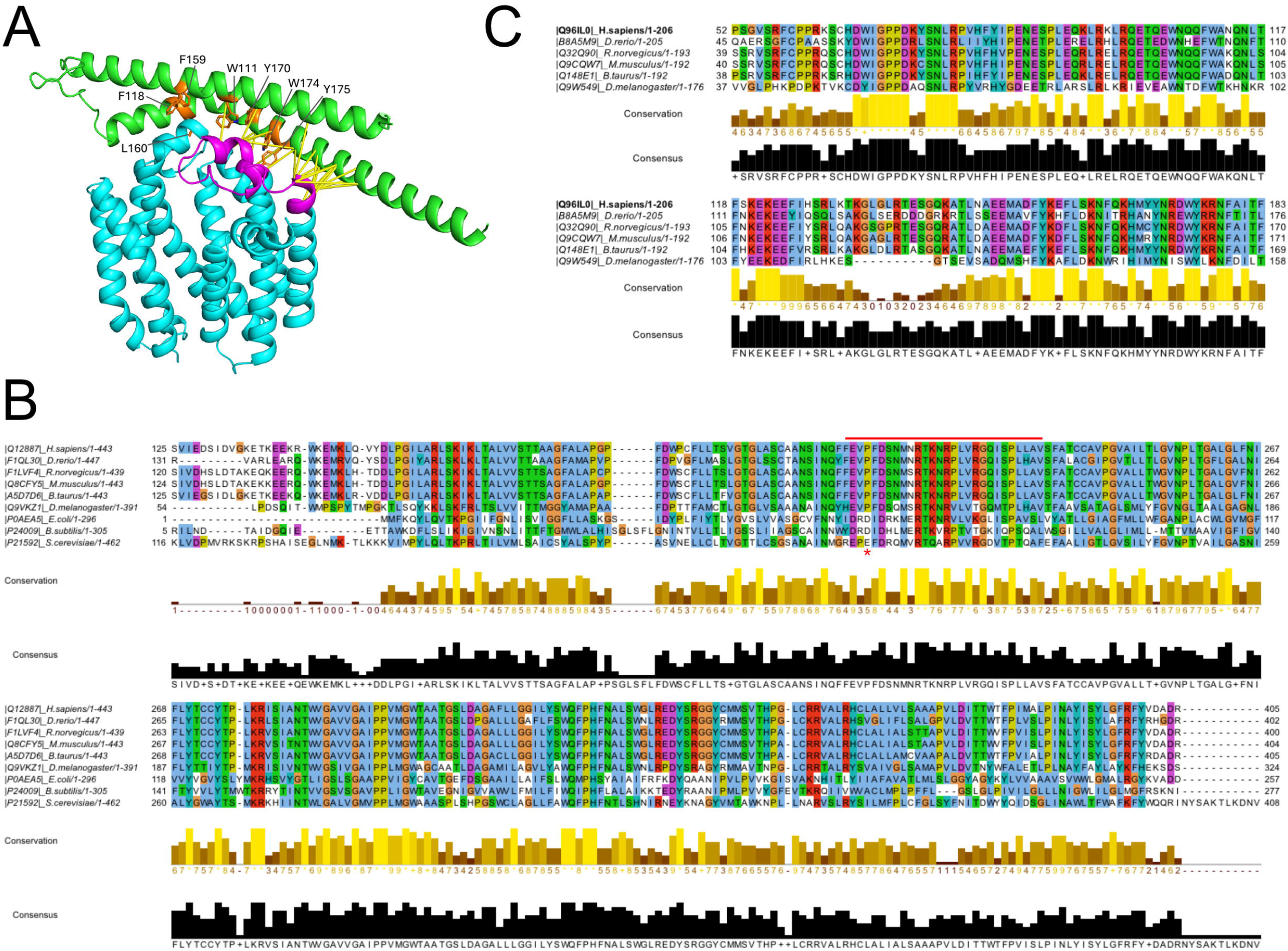
COA8 is predicted to interact with the COX10 central cavity catalytic domain through a set of conserved residues. (**A**) The AlphaFold model of the COA8-COX10 complex. COA8 and COX10 are colored green and cyan, respectively. The helix loop region (HL2-3) in between the second and third transmembrane segments of COX10 is colored magenta. Large hydrophobic residues of COA8 in the interface are colored orange and labeled. Yellow sticks are shown between residue pairs that have high intermolecular contact probability (above 0.9) according to AlphaFold. **(B)** Sequence conservation and alignment of COX10 throughout evolution. Alignments have been performed with Clustal Omega and analyzed with Jalview 2.11.3.2. The central pocket is highlighted with a red bar on top of the sequence and the proline residue (P213) is highlighted with a red asterisk. **(C)** Sequence conservation and alignment of COA8 throughout evolution in metazoans. Alignments have been performed with Clustal Omega and analyzed with Jalview 2.11.3.2.

**Table 1.**
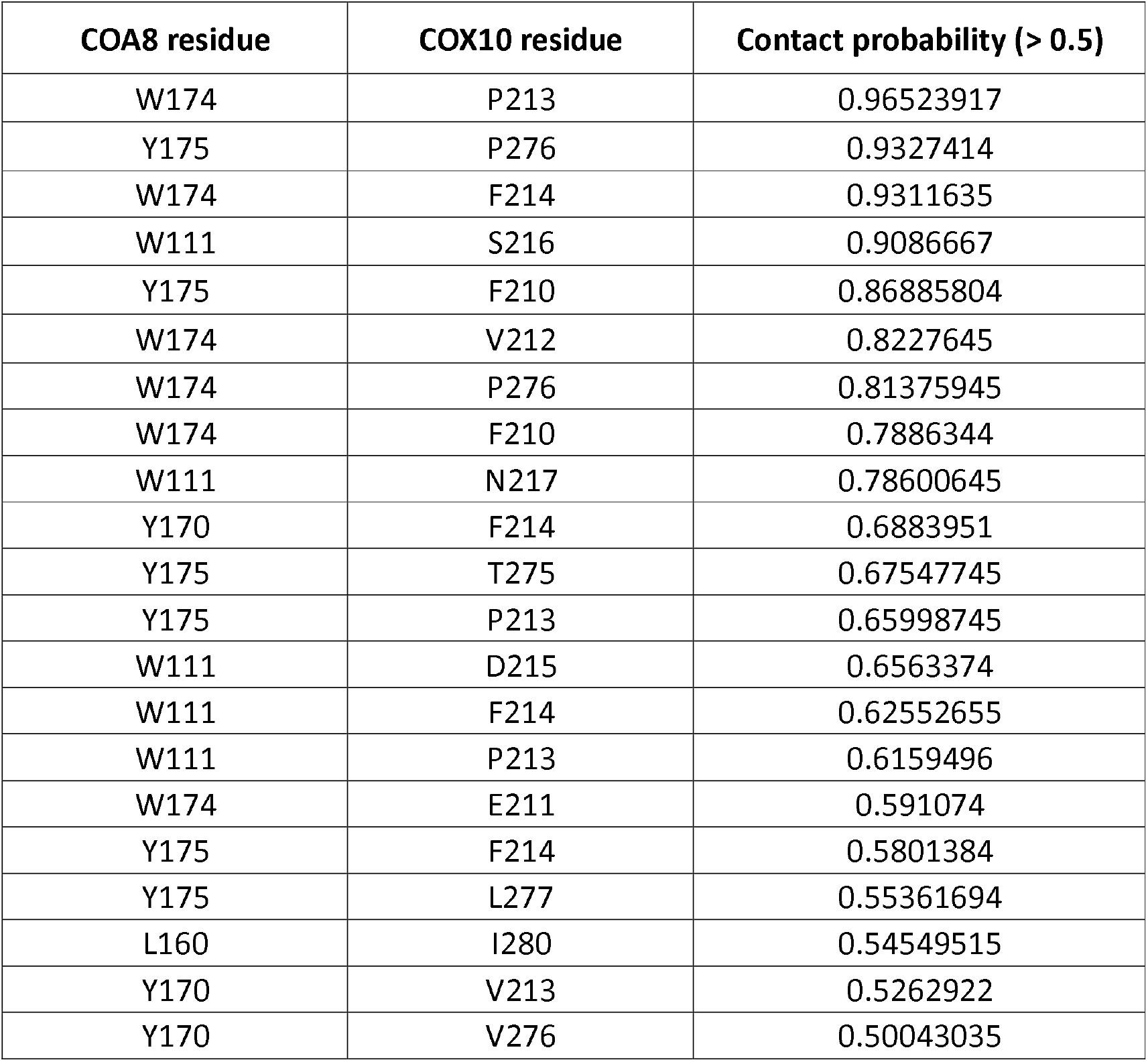
Relevant residues for the predicted interactions between COA8 and COX10.

### A COA8 W174A variant is detrimental for function, but does not prevent the interaction with COX10

To determine the role of the predicted relevant aromatic residues in COA8, we generated mutant versions of the GFP-tagged COA8 and transduced the S6 immortalized fibroblast cells with different variants. Lentiviral vectors containing the WT version of COA8^GFP^, as well as the GFP alone, were transduced again in parallel to the mutated sequences, to serve as positive and negative controls, respectively. Of the potentially relevant interacting amino acids (see above), we chose one residue in each of the three positions of the COA8 domain (beginning, middle and end). The F118 residue was especially interesting for this study, since a homozygous F118S variant was found in a patient suffering of severe leukoencephalopatic disease (Melchionda *et al*., 2014). Thus, the COA8 mutations generated and tested were: F118A, F118S, F159A and W174A. As we had previously observed (Signes *et al*., 2019), expression of WT COA8^GFP^ in the patient-derived fibroblasts was able to fully restore MT-CO2 protein levels (**Figure 4A**) and cytochrome *c* oxidase assembly and activity (**Figure 4B** and **Figure 4C**). Expression of the F118A^GFP^, F118S^GFP^ and F159A^GFP^ variants rescued the cIV amounts and activity to a similar extent as the WT-COA8^GFP^ (**Figure 4A** and **4B**). Contrariwise, the S6 cells expressing the W174A^GFP^ variant continued to show profound cIV deficiency (**Figure 4A, 4B** and **4C**), despite that the COX10 levels were recovered (**Figure 4A**). This is most likely due to the preserved interaction between the W174A^GFP^ variant and COX10, as demonstrated by the fact that they still co-immunoprecipitate (**Figure 4D**). Considering that COA8 is most probably binding to the catalytic domain of COX10, we hypothesized that COA8 is a factor that stabilizes and modulates COX10 activity. Since heme A levels are reduced in patient-derived fibroblasts carrying pathological mutations in *COX10* (Antonicka *et al*., 2003a), we measured the amount of heme A and heme B present in our cells. The relative amounts of heme A to heme B were decreased about three-fold in the S6+GFP cells compared with the complemented S6+COA8^GFP^ (**Figure 4E**). Consistent with the detrimental effects of the W174A COA8 variant on cIV stability/assembly and activity, heme A levels were also decreased in the S6+W174^GFP^ cells to about half of the S6+GFP negative control and 6-fold compared with the positive control S6+COA8^GFP^ cells.

**Figure 4.**
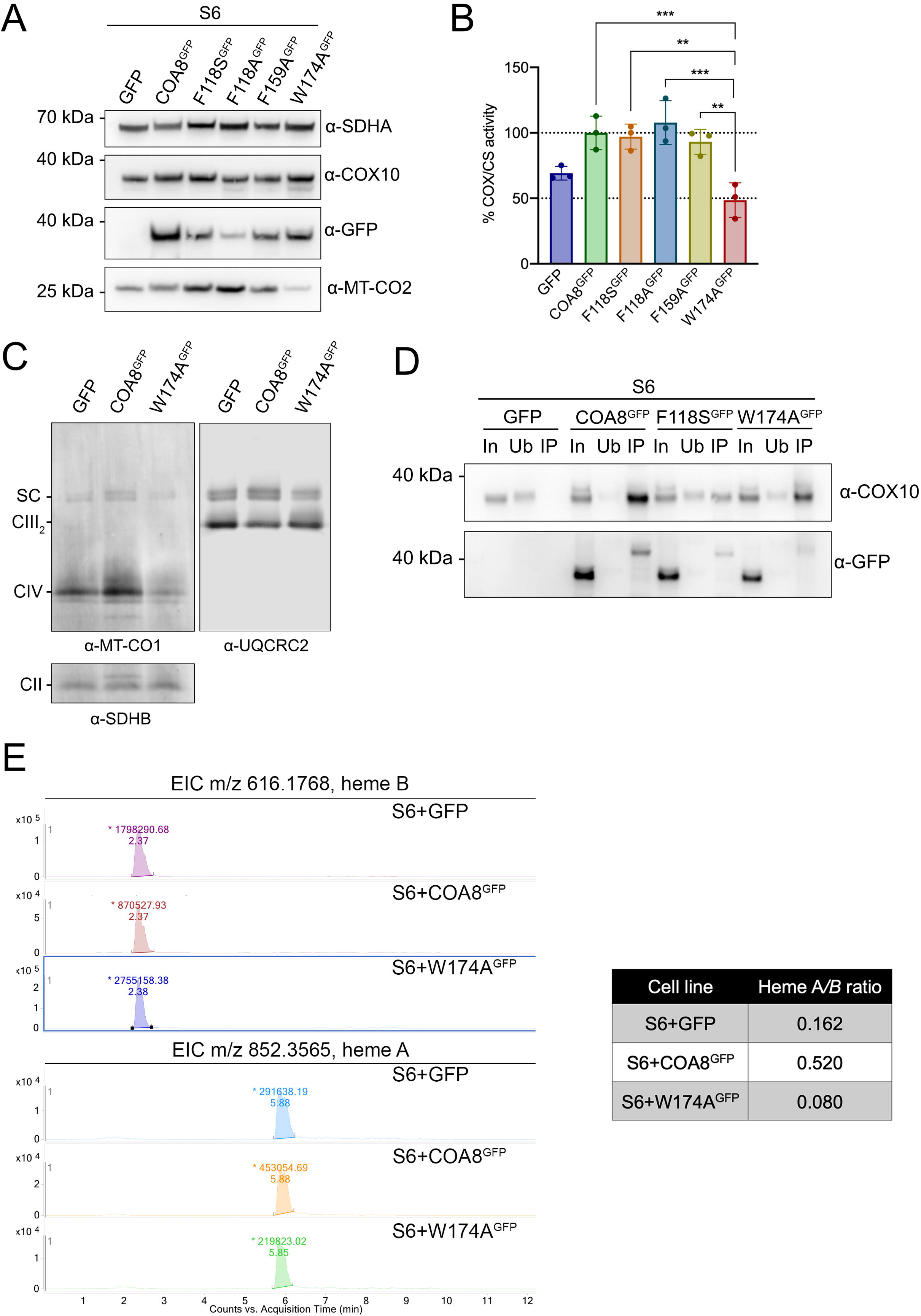
A COA8 W174A variant is detrimental for function, but does not prevent the interaction with COX10. (**A**) Steady-state levels of GFP, MT-CO2, COX10 and SDHA in uncomplemented patients’ fibroblasts (GFP) and complemented with wild-type COA8^GFP^ or GFP-tagged COA8 variants F118S, F118A, F159A and W174A. **(B)** Kinetic enzyme activity of complex IV (COX), normalized by citrate synthase (CS) activity, in mitochondria-enriched fractions from uncomplemented patient fibroblasts (GFP) and complemented with wild-type COA8-GFP or mutant variants F118S, F118A, F159A and W174A. Data are plotted as mean ± S.D of the normalized activities (n = 3 biological replicates, one-way ANOVA with Dunnett’s multiple comparisons, **p ≤ 0.01, ***p ≤ 0.001). **(C)** BN-PAGE and western blot immunodetection of MRC complexes in DDM-solubilized mitochondria-enriched fractions from uncomplemented patients’ fibroblasts (GFP) and complemented with wild-type COA8-GFP or the mutant variant W174A using antibodies against MT-CO1, UQCRC2 and SDHB. CII = complex II; CIV = monomeric complex IV; CIII_2_ = dimeric complex III; SC = supercomplex III_2_+IV **(D)** Western blot analysis of anti-GFP affinity purified complexes from uncomplemented patient S6 fibroblasts (GFP) and complemented with wild-type COA8^GFP^ or the mutant variant W174A^GFP^. In = total mitochondrial-enriched fractions (input); Ub = unbound material; IP = immunoprecipitated material. Samples were probed with antibodies against GFP and COX10. **(E)** Extracted ion chromatograms (EIC) for heme B (616.1768) and heme A (m/z 852.3565) obtained for each of the three isolated mitochondria samples from S6+GFP, S6+COA8^GFP^ and S6+W174A^GFP^ cells. The upper numbers on top of each peak correspond to the area under the curve values, and the lower number is the acquisition time in minutes. The table shows the values of heme A to heme B ratios, calculated dividing the area under the curve for heme A by the area under the curve for heme B obtained for each sample.

### COA8 loss-of-function blocks MT-CO1 metallochaperone-mediated maturation and assembly into supercomplexes

In order to investigate in detail the structural/functional defects induced by the absence of COA8 as well as by the W174A COA8 variant, we carried out complexome profiling (Cabrera-Orefice *et al*, 2021) to analyze the assembly of the proteins involved in cIV biogenesis in the S6+GFP, S6+COA8^GFP^ and S6+W174A^GFP^ cells. COA8 loss-of-function, either by complete absence of the protein or caused by the W174A variant, produced an accumulation of the early cIV assembly module (COX4I1+COX5A+HIGD1A) and of the MT-CO1 subunit (**Figure 5A**). This was linked to decreased incorporation of cIV subunits into supercomplexes SCIII_2_+IV and SCI+III_2_+IV (respirasomes), both relative to the amount of monomeric cIV and in comparison with the control (S6+COA8^GFP^) (**Figure 5A**). The lower amounts of cIV-containing SCs are not due to a decrease in the amounts of complexes I and III_2_, which were in fact more abundant in COA8-deficient cells, especially in the S6+W174A^GFP^, with an enhanced formation of SCI+III_2_ (**Supplemental Figure S2**). A similar phenomenon was observed in *Coa8-*deficient *Drosophila melanogaster* mitochondria (Brischigliaro *et al*., 2023).

**Figure 5.**
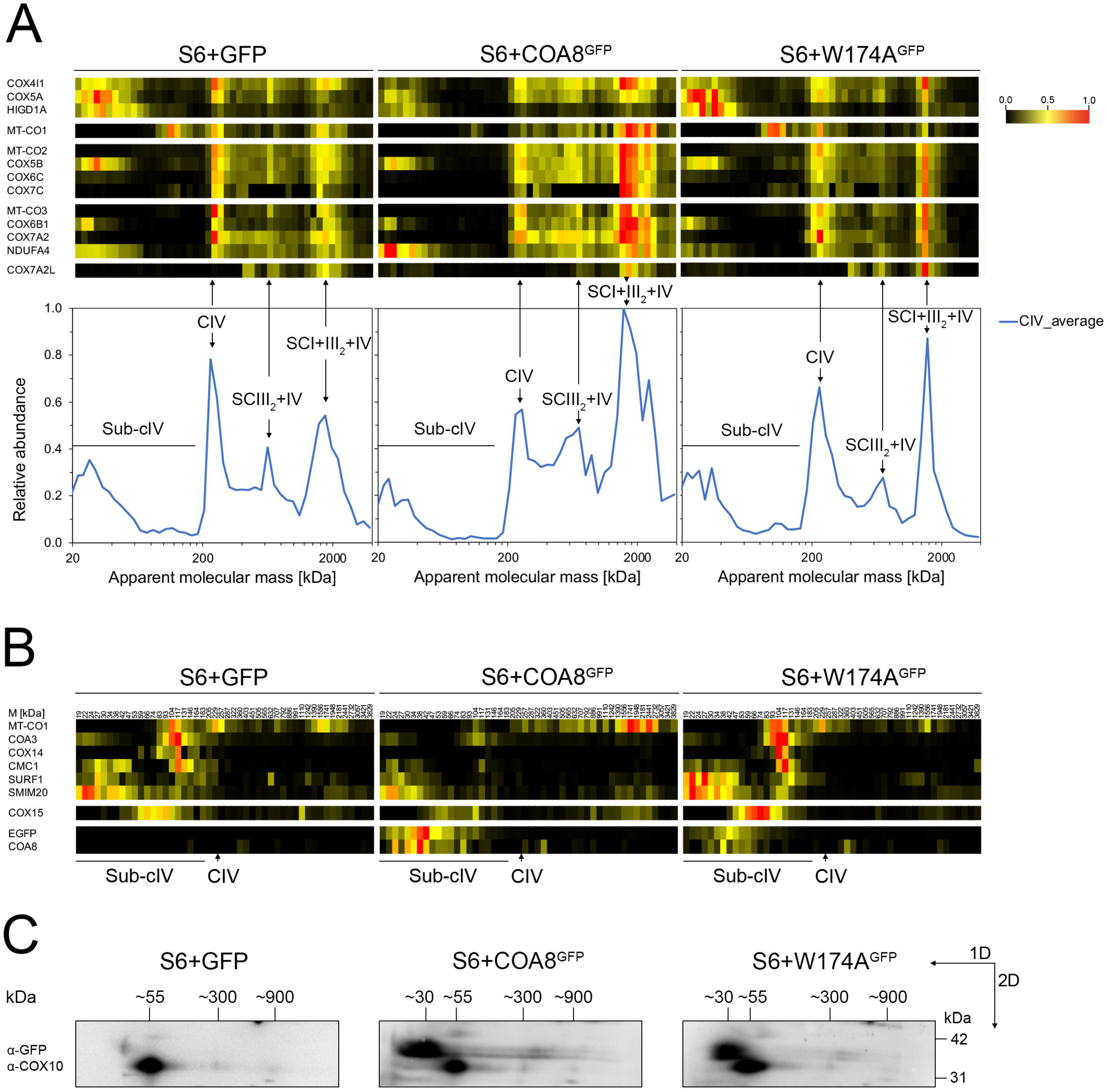
COA8 loss-of-function blocks MT-CO1 metallochaperone-mediated maturation and assembly into supercomplexes. (**A**) Heatmaps of peptide relative intensity along the BN-PAGE lanes, corresponding to the cIV structural subunits in each of the analyzed cell lines (S6+GFP, S6+COA8^GFP^, S6+W174A^GFP^), divided into groups according to the assembly module they belong to (see Figure 1C). The peptide intensity heatmap of COX7A2L (aka SCAFI), which binds to SCIII_2_+IV and to a proportion of SCI+III_2_+IV, as well as to cIII_2_ (Fernández-Vizarra *et al*, 2022; Perez-Perez *et al*, 2016; Vercellino & Sazanov, 2021), is shown in order to locate the position of the SCs within the heatmaps. The profiles correspond to the averaged relative peptide intensities of the detected cIV subunits. **(B)** Heatmaps of peptide relative intensity along the BN-PAGE lanes, corresponding to MT-CO1 and the binding chaperones that mediate its maturation, also known as the ‘MITRAC’ complex (Dennerlein *et al*., 2023; Timon-Gomez *et al*., 2018), as well as those of COX15 (heme A synthase) and COA8 and EGFP. **(C)** Immunodetection of COA8^GFP^ and W174A^GFP^ (using an anti-GFP antibody) and of COX10, in western blot membranes of two-dimensional BN-PAGE. Complexes were separated first (1D) in a 4%->16% gradient native gel and then in a denaturing 4%->12% gradient SDS-PAGE (2D).

It is still unclear how the heme A moieties are incorporated into the MT-CO1 apoprotein, but it must occur shortly after the subunit is translated at the vicinity of the IMM and while it is still bound to a number of stabilizing chaperones: COA3, COX14, CMC1, SURF1 and SMIM20 (Dennerlein *et al*., 2023; Timon-Gomez *et al*., 2018). This set of proteins has also been denominated the “<u>mi</u>tochondrial <u>t</u>ranslation <u>r</u>egulation <u>a</u>ssembly intermediate of <u>c</u>ytochrome *c* oxidase” (MITRAC) (Mick *et al*, 2012). In the S6+GFP, all of these chaperones were accumulated and, for the most part, co-migrated with MT-CO1 at a calculated molecular mass of 93-117 kDa (**Figure 5B**). In the complemented cell line with WT COA8, the accumulation of MT-CO1 chaperones was much lower, compatible with a progression in cIV assembly, with significantly less stalled MT-CO1. In the S6 cells expressing the W174A^GFP^ variant, we observed a similar stalling of MT-CO1 and accumulation of chaperones as in the null-mutant cells (S6+GFP). This observation indicates that the defect induced by the W174A mutation is also at the level of MT-CO1 maturation/metalation. The majority of both COA8^GFP^ and W174A^GFP^ migrated to lower molecular masses, estimated to be between 30 and 42 kDa according to its migration on Blue Native (BN)-PAGE (**Figure 5B** and **Figure 5C**), with a slight overlap with the COX10 signal (**Figure 5C**). COX10 peptides escaped identification by MS in the complexome profiling analysis, but we were able to immunodetect it by Western blot of a denaturing second dimension (2D) after a first BN-PAGE dimension (1D) (**Figure 5C** and **Supplemental Figure S3**). Most of the COX10 signal was detected at an apparent molecular mass of ∼50 kDa with two additional weaker signals at ∼270 and ∼900 kDa. Remarkably, there were no noticeable differences in the COX10 migration pattern among the three cell lines. The reason for this might be that only very low levels of COA8 are necessary to carry out its function (Signes *et al*., 2019). Peptides corresponding to the other enzyme of the heme A synthesis pathway (COX15), were detected by MS mostly migrating between ∼53 kDa and ∼104 kDa in the BN-PAGE, indicating that the protein could be in homodimeric form or interacting with another protein of similar size (**Figure 5B**). Similarly to the MT-CO1 chaperones, COX15 was accumulated more in the two COA8-deficient cells than in the complemented fibroblasts. Lower amounts of COX15 appeared bound to high molecular mass complexes, which were different in each one of the three cell lines. The reasons for this could be due to COX15 being able to also bind cIII_2_, as described in yeast (Herwaldt *et al*, 2018).

Recently, the existence of a metallochaperone complex that coordinates the formation of the cIV active centers during the early steps of MT-CO2 and MT-CO1 assembly has been described. In this scenario, the copper chaperone COX11 appears to play a role in the formation of both the Cu_A_ and Cu_B_ centers, being the copper atom in the Cu_B_ center inserted concomitantly to Heme a3 and after Heme a (Nyvltova *et al*., 2022). During the maturation process, and to continue the assembly of cIV, the chaperones must be released from MT-CO1 and MT-CO2 at different steps of the process (Timon-Gomez *et al*., 2018). The main components of the copper metallochaperone complex, i.e., SCO1, SCO2, COA6, COX16 and COX11 were also accumulated at molecular masses lower than 200 kDa in S6+GFP and S6+W174A^GFP^, when compared with the control expressing WT COA8^GFP^ (**Figure 6A**). The main difference was found in the levels and migration pattern of COX11 in the COA8-deficient cells, having the highest peptide intensities at ∼47-53 KDa vs. ∼34 kDa in the control. In addition, SCO1 was markedly accumulated in the same 53 kDa peak in the S6+GFP cells (**Figure 6A**). The migration behavior of SCO2 was also different in the cells expressing the functional COA8^GFP^, in comparison with the two defective cell lines (**Figure 6A**).

**Figure 6.**
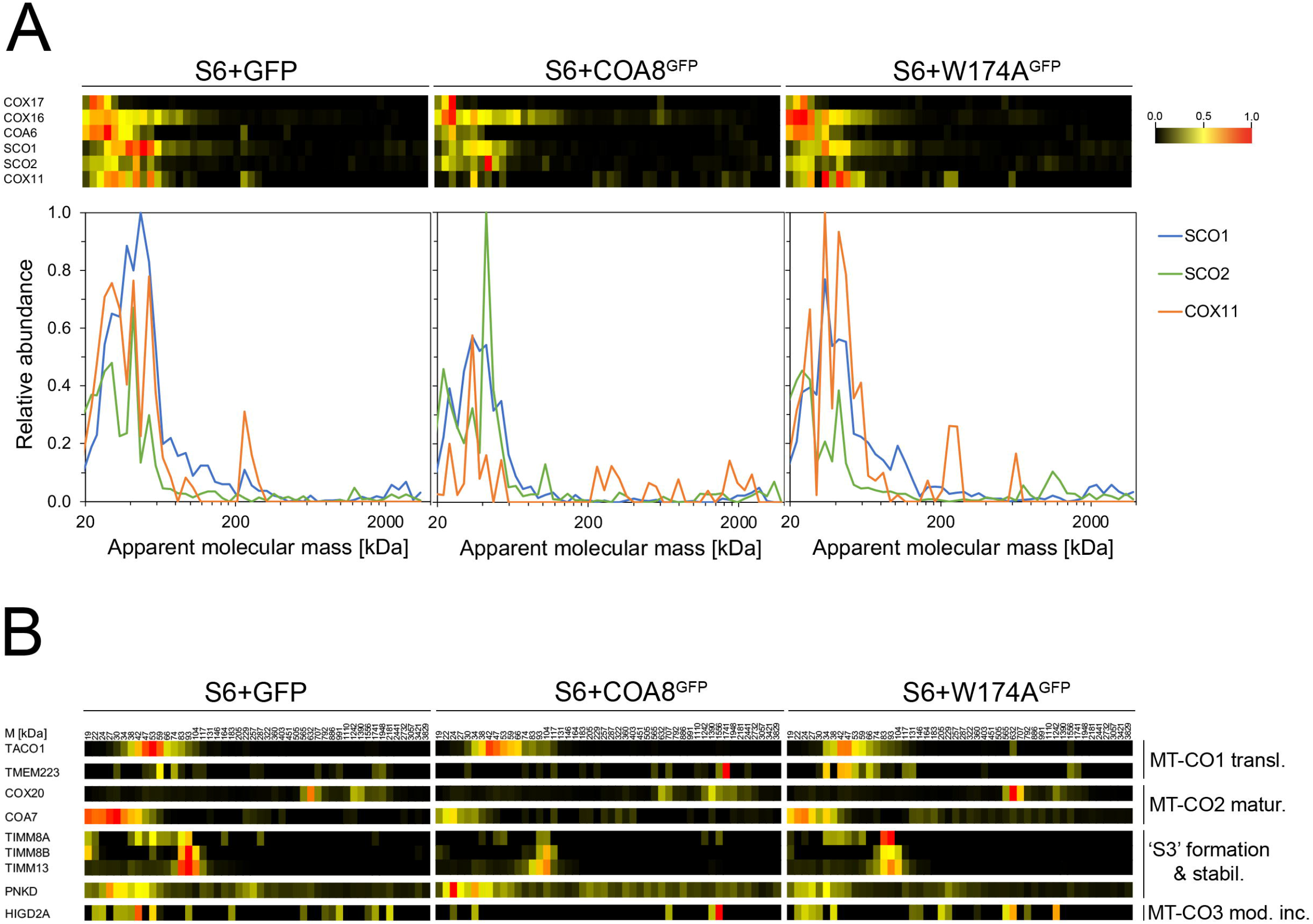
Alterations in the copper metallochaperones due to COA8-loss-of-function. (**A**) Heatmaps of peptide relative intensity along the BN-PAGE lanes, corresponding to the proteins involved in Cu delivery during cIV assembly in each of the analyzed cell lines (S6+GFP, S6+COA8^GFP^, S6+W174A^GFP^). The profiles corresponding to the proteins which were the most changed between the control (S6+COA8^GFP^) and the COA8-deficient cells (S6+GFP, S6+W174A^GFP^), i.e., SCO1, SCO2 and COX11, are shown for comparison. **(B)** Heatmaps of peptide relative intensity along the BN-PAGE lanes, corresponding to different proteins involved in cIV assembly, divided into groups according to the step they are involved in (MT-CO1 translation; MT-CO2 maturation; ‘S3’ subcomplex formation and/or stabilization; MT-CO3 module incorporation).

Other cIV assembly factors acting either upstream or downstream of MT-CO1 chaperone-mediated maturation (**Figure 1C**) accumulated in S6+GFP and S6+W174A^GFP^ cells (**Figure 6B**). For example, peptides corresponding to TMEM223, which is a mitoribosome-interacting early cIV assembly factor that also binds the early ‘MITRAC’ complex (Dennerlein *et al*, 2021), showed higher intensities at lower molecular mass positions in the COA8-deficient cells, whereas in the control it was mainly detected at a position corresponding to ∼1741 kDa, which was the same as the mitochondrial ribosome (**Figure 6B**, **Supplemental Table S4**). The migration patterns and abundance of factors involved in the maturation of MT-CO2, such as COX20 (Bourens *et al*, 2014) or COA7 (Formosa *et al*, 2022), as well as in the formation and/or stabilization of the ‘S3’ subcomplex, such as TIMM8A, TIMM8B, and TIMM13 (Anderson *et al*, 2023) and MR-1S (PNKD) (Vidoni *et al*., 2017), were altered in the COA8-deficient cells. Finally, HIGD2A, the factor facilitating the last step of cIV assembly, i.e. the incorporation of the MT-CO3 module (Hock *et al*, 2020; Timon-Gomez *et al*., 2020), also accumulated at lower molecular masses in the two uncomplemented S6 cell lines (**Figure 6B**).

Altogether, these observations consistently indicate a blockage in the metallochaperone-mediated maturation of MT-CO1 when COA8 is either absent or non-functional.

## DISCUSSION

Despite recent significant advances, there are still many open questions concerning the steps and players necessary to achieve the proper assembly, organization and function of the respiratory chain (Fernandez-Vizarra & Ugalde, 2022; Signes & Fernandez-Vizarra, 2018). This is, however, a fundamental question not only for mitochondrial biology and bioenergetics, but also for medicine, because defects in the assembly of OXPHOS lead to the development of disease in humans (Fernandez-Vizarra & Zeviani, 2021b). CIV assembly is a highly intricate and regulated process, where the number of assembly factors involved duplicates the number of actual structural components (Timon-Gomez *et al*., 2018). Pathological variants in the genes encoding many of these assembly proteins are the cause of mitochondrial disease associated with cIV deficiency (Brischigliaro & Zeviani, 2021). In this work, we aimed to get a deeper understanding of the function of COA8, part of the group of mitochondrial disease-associated cIV assembly factors. Even though the protein was originally named as APOPT1, work done by our group clearly showed that this protein is associated to the IMM and facing the matrix, and that it plays a role in the assembly of cIV in humans, mouse and *Drosophila melanogaster* (Brischigliaro *et al*., 2023; Brischigliaro *et al*., 2019; Melchionda *et al*., 2014; Signes *et al*., 2019). Thus, the name was officially changed to Cytochrome *c* Oxidase Assembly factor 8 (COA8). However, the exact molecular role played by COA8 within the cIV assembly pathway was still largely unknown. In this study, we have been able to experimentally prove the interaction between COX10 and COA8, the latter modulating the function of the former. In addition, thorough proteomic and functional analyses have clearly demonstrated that COA8 loss-of-function leads to impaired MT-CO1 maturation, thus preventing the normal progression of cIV assembly. Since cIV is the sole heme A-containing enzyme and because of the potential toxicity of free heme A, and of the absence of intra-mitochondrial heme degradation pathways, heme A synthesis is likely to be strictly coupled to cIV assembly (Khalimonchuk *et al*, 2012). In addition, the synthesis of heme O from heme B, catalyzed by COX10 in mitochondria, appears to be the rate-limiting step in heme A formation (Khalimonchuk *et al*., 2012; Wang *et al*, 2009). Therefore, the block in MT-CO1 maturation and impaired cIV assembly observed in subjects carrying loss-of-function variants in *COA8*, and in *Coa8^KO^* animal models, is most likely due to decreased heme A availability because of diminished catalytic activity of COX10. Accordingly, heme A levels were drastically reduced in the absence of COA8 function, compared with the control cells. The finding that in human mitochondria COA8 binds to COX10 is extremely intriguing, since heme O synthases are found in many prokaryotic organisms and, therefore, appeared very early during evolution (Rivett *et al*., 2021). In contrast, COA8 appeared much later in evolution, being only present in metazoan species. Therefore, it is surprising that the proper stability and function of COX10 in metazoans clearly relies on the presence of a functional COA8, as we have here demonstrated. COA8 is confidently predicted to bind to the catalytic domain of COX10 (Pei *et al*., 2022), which is a loop between transmembrane helices 2 and 3, facing the mitochondrial matrix. This location coincides with the experimentally determined topology of COA8 (Signes *et al*., 2019). The COX10 catalytic domain contains several residues that are highly conserved from bacteria to mitochondria, and essential for the heme O synthase catalytic activity (Rivett *et al*., 2021). Interestingly, when looking at the sequence alignments, the catalytically relevant residues (highlighted in **Figure 3B**) are highly conserved from bacteria to humans except for *E. coli* D63. This position is occupied by P213 in human COX10 and this particular Pro residue is conserved across metazoans. In fact, P213 was predicted with high confidence to be one of the top residues interacting with COA8, mainly with COA8 W174 (**Table 1**), which has been proven to be necessary for function. Pro-Trp interactions might play an important conformational and stabilization role in proteins (Biedermannova *et al*, 2008). Therefore, the interaction between COX10 P213 and COA8 W174 is most likely to be important to keep the catalytic loop of COX10 in an active conformation. However, if this position is occupied by an acidic amino acid, like in bacteria or yeast, the interaction with an extra factor might not be necessary. This is a possibility that will be experimentally addressed in the future. In fact, yeast COX10 appears to form homo-oligomeric complexes of a size of ∼250-300 kDa and this oligomerization is mediated by the Coa2 assembly factor (Khalimonchuk *et al*., 2012), which does not have a human homolog. In human mitochondria, we have observed that most of COX10 is found in monomeric form, with only residual amounts incorporated in higher order complexes. Therefore, the way COX10 is stabilized varies between *S. cerevisiae*, where no COA8-like protein exists, and human mitochondria, in which COA8 is present.

Further, we have observed that COA8 loss-of-function globally affects the process of cIV metalation. This is illustrated by the fact that the factors involved, which are part of the denominated ‘metallochaperone complex’, are accumulated in the deficient patient-derived cells. The patterns of this accumulation are very similar between the S6 cells expressing only GFP (negative control) and those expressing the W174A-COA8^GFP^ variant. This is yet another confirmation that this mutation, even if the protein is stable and able to interact with COX10, induces a defect at the same functional level, i.e., heme A synthesis/incorporation. Moreover, our complexome profiling data revealed noticeable changes in the migration pattern of SCO1 and COX11 between the COA8-functional and dysfunctional mitochondria, being found to accumulate at higher molecular mass positions in the latter. SCO1 and COX11 contain redox-sensitive Cys residues necessary for their copper chaperone function (Povea-Cabello *et al*, 2024). In addition, COX11 forms a homodimer in which the Cys residues are likely to be involved in Cu coordination between the two monomers (Carr *et al*, 2002; Nyvltova *et al*., 2022). It is possible that the higher apparent molecular mass of COX11 in the COA8-deficient cells is due to the formation of the dimer, and that more COX11 is found in dimeric form than in monomeric form due to a defect in Cu delivery, which seems to be linked to MT-CO1 hemylation (Nyvltova *et al*., 2022). Interestingly, the redox state of the Cys in SCO1 and COX11 are changed when COX10 expression is downregulated (Nyvltova *et al*., 2022), providing another link, and additional evidence, to the involvement of COX10 dysfunction in our models, caused by COA8 impairment.

In conclusion, the present work provides novel insights into the pathophysiological role of COA8, providing the molecular basis underlying mitochondrial disease associated with loss-of-function of this factor, as well as elucidating its mechanistic role in the biogenesis of metazoan mitochondrial cIV.

## ACKNOWLEDGEMENTS

We would like to thank Jana Meisternknecht (Goethe University Frankfurt, Germany) for excellent technical assistance. This research was funded by Telethon Foundation-Cariplo Foundation Alliance (grant GJC21014 to E.F.-V.), AFM-Téléthon (24962 to E.F.-V.), Department of Biomedical Sciences—UNIPD (FERN_FAR22_01 to E.F.-V.), Telethon Foundation (GGP20013 to C.V.), AFM-Téléthon (23706 to C.V.), Associazione Luigi Comini Onlus (MitoFight2, to C.V.), the Department of Biomedical Sciences—UNIPD (SID2022-VISC_BIRD2222_01 to C.V.), the Netherlands Organization for Health Research and Development (ZonMW TOP 91217009 to S.A.) and the Deutsche Forschungsgemeinschaft (DFG) (SFB1531 (S01, project number 456687919) to A.C.-O.). K.C. is the recipient of an EMBO Postdoctoral Fellowship (ALTF 710-2022).

## AUTHOR CONTRIBUTIONS

Conceptualization, M.B. and E.F.-V.; Methodology, M.B. and E.F.-V.; Software, A.C.O., J.P., Q.C.; Validation, M.B., K.C., A.C.O., J.P., C.F., M.R., S.P.C., S.B., G.A., S.A., Q.C., M.Z., C.V. and E.F.-V.; Formal analysis, M.B., K.C., A.C.O., J.P., M.R., G.A., Q.C. and E.F.-V.; Investigation, M.B., K.C., A.C.O., J.P., C.F., M.R., S.P.C. and E.F.-V.; Resources A.C.O., J.P., S.P.C., S.B., G.A., S.A., Q.C., M.Z., C.V. and E.F.-V.; Data Curation M.B., K.C., A.C.O., J.P., C.F., G.A., S.A., Q.C. and E.F.-V.; Writing-Original Draft M.B. and E.F.-V.; Writing-Review & Editing M.B., K.C., A.C.O., J.P., C.F., M.R., S.P.C., S.B., G.A., S.A., Q.C., M.Z., C.V. and E.F.-V.; Visualization M.B., K.C., A.C.O. and E.F.-V.; Supervision S.B., G.A., S.A., Q.C., M.Z., C.V. and E.F.-V.; Project administration S.B., G.A., S.A., Q.C., M.Z., C.V. and E.F.-V.; Funding acquisition C.V. and E.F.-V.

## DATA AVAILABILITY

The datasets and computer code produced in this study will be made available upon publication in the following databases:

– Protein MS data: PRIDE (https://www.ebi.ac.uk/pride/)
– Complexome profiling data: CEDAR (van Strien *et al*., 2021) (https://www3.cmbi.umcn.nl/cedar/browse/)

**Supplemental Figure S1. Related to Figure 1.**
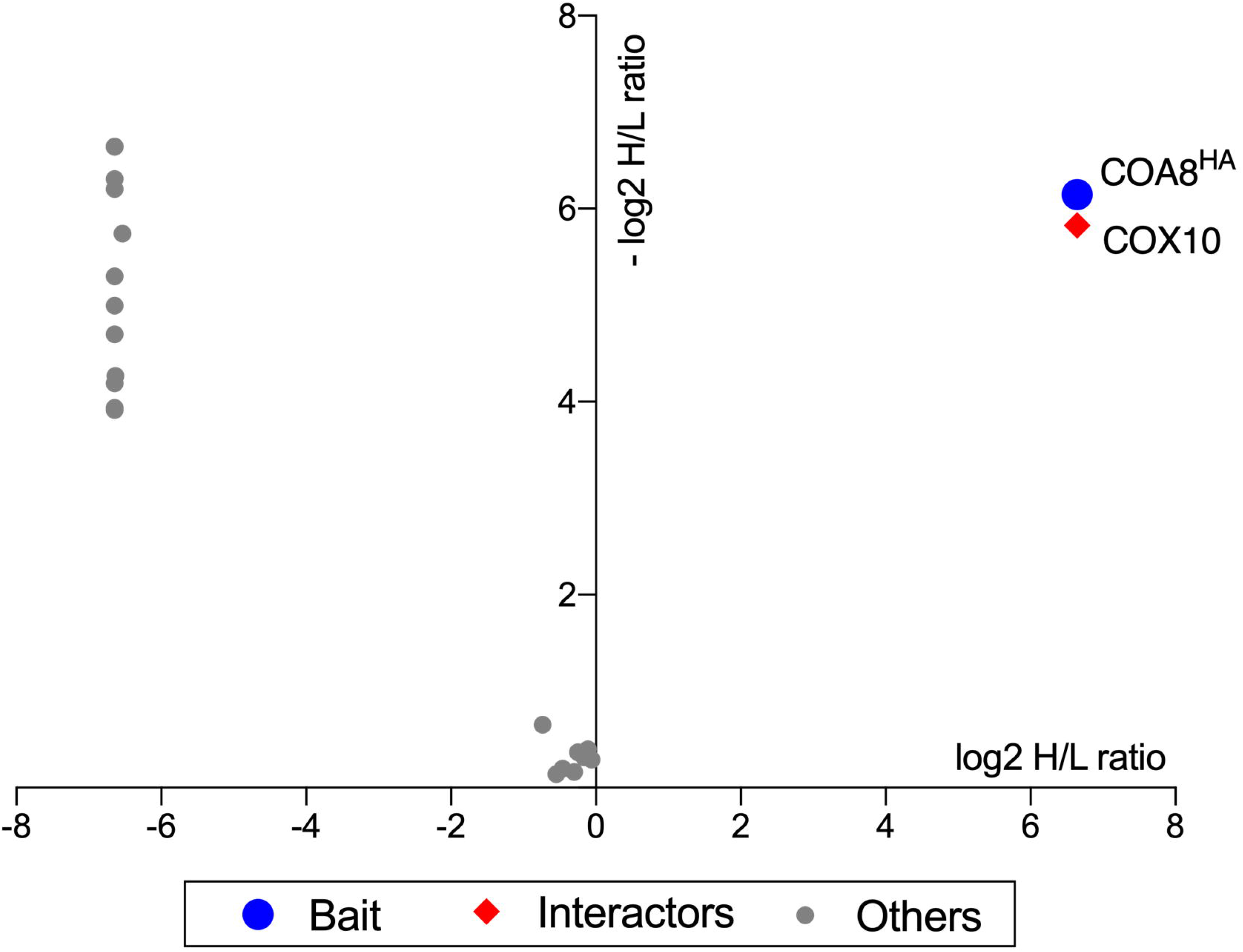
Reverse SILAC-labeled mitochondria-enriched fractions from 143B cell lines were isolated, mixed, and subjected to anti-HA immunocapture after 1h treatment with 100 μM H_2_O_2_. The values in the x axis correspond to the log2 heavy-to-light (H/L) ratio of the proteins detected in experiment 1, where the heavy (H)-labeled COA8-HA expressing cells and unlabeled (L) GFP expressing cells were mixed. The values in the y axis correspond to the inverted log2 H/L ratio (−log2 H/L) of the proteins detected in experiment 2, where the unlabeled (L) COA8-HA expressing cells and labeled (H) GFP expressing cells were mixed. Blue dot = bait (HA); red squares = interactors; grey dots = others.

**Supplemental Figure S2. Related to Figure 5.**
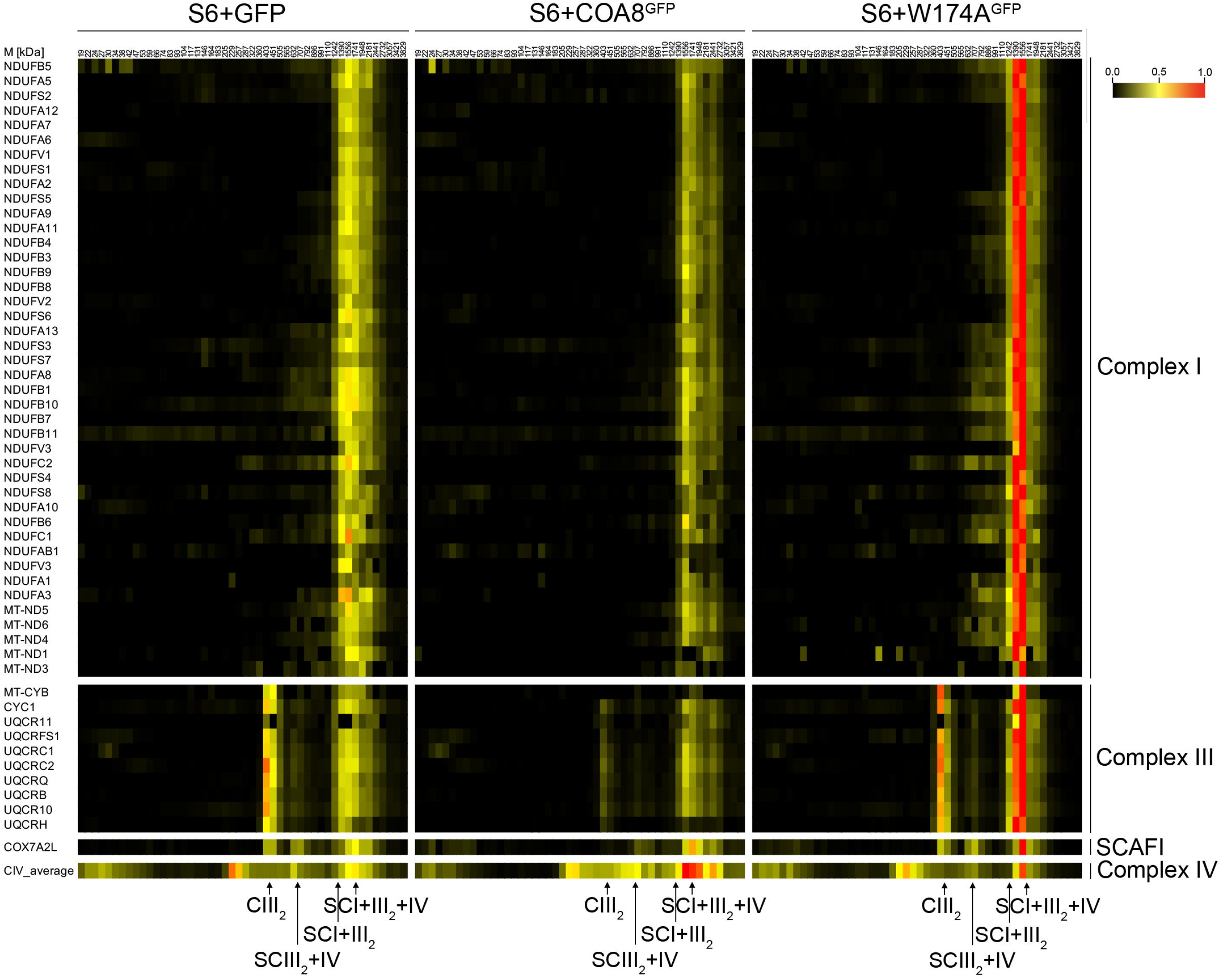
Heatmaps of peptide relative intensity along the BN-PAGE lanes, corresponding to the structural subunits of cI and cIII_2_ in each of the analyzed cell lines (S6+GFP, S6+COA8^GFP^, S6+W174A^GFP^). The peptide intensity heatmap of COX7A2L (aka SCAFI), which binds to SCIII_2_+IV and to a proportion of SCI+III_2_+IV, as well as to cIII_2_ (Fernández-Vizarra *et al*., 2022; Perez-Perez *et al*., 2016; Vercellino & Sazanov, 2021) and of the average of the cIV structural subunits, are shown in order to locate the position of the SCs within the heatmaps.

**Supplemental Figure S3. Related to Figure 5.**
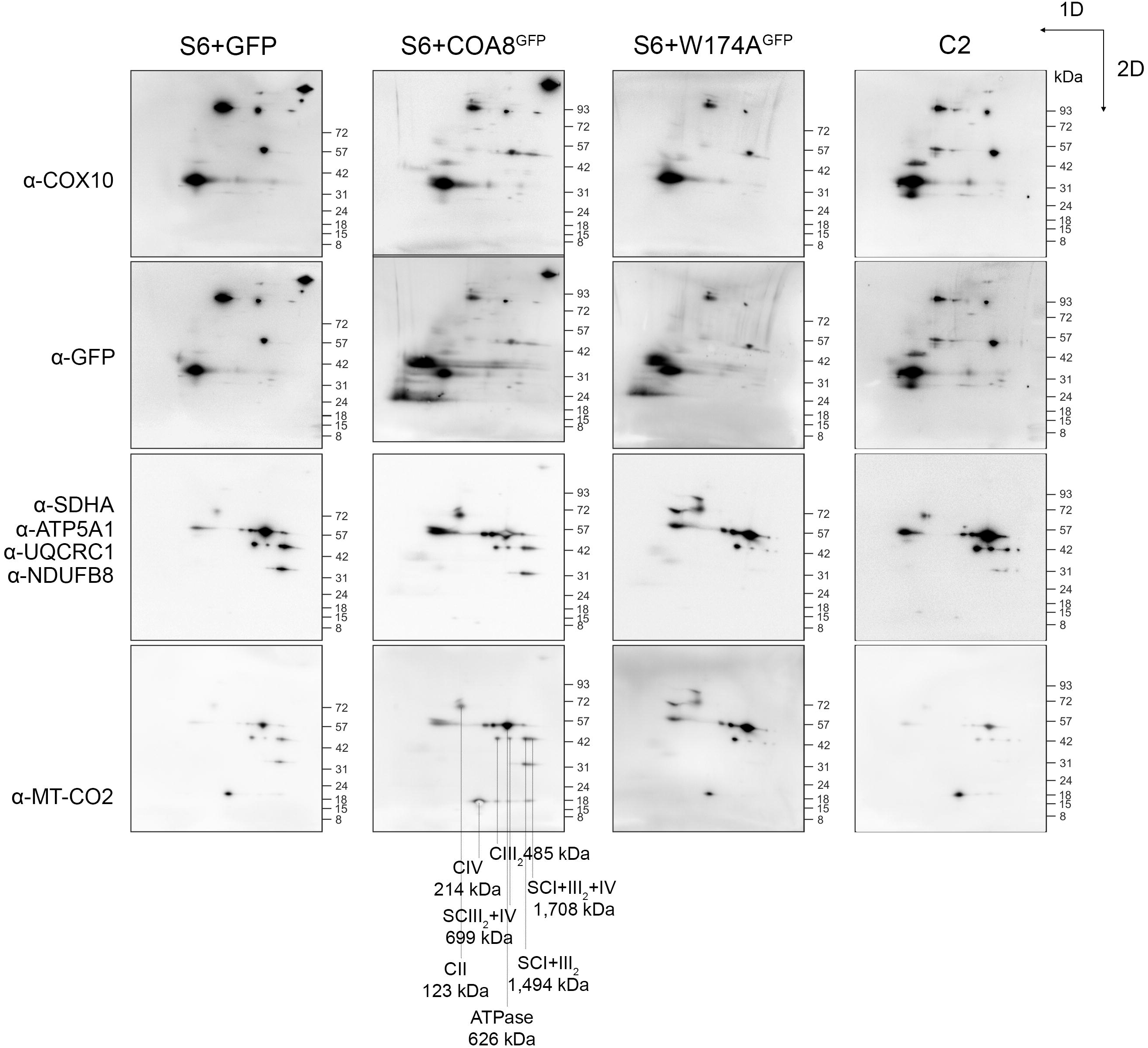
Immunodetection of COA8^GFP^ and W174A^GFP^ (using an anti-GFP antibody), COX10 and the OXPHOS complexes, using antibodies against one subunit of each complex (cI: NDUFB8, cII: SDHA; cIII_2_: UQCRC1; cIV: MT-CO2 and ATPase: ATP5A1) in Western blot membranes of two-dimensional BN-PAGE. Complexes were separated first (1D) in a 4%->16% gradient native gel and then in a denaturing 4%->12% gradient SDS-PAGE (2D). The molecular masses of each complex used to estimate the molecular sizes of the COX10-containing complexes, calculated by measuring the migration distances in the BN-PAGE, are indicated at the bottom.

**Supplemental Table S1.** Processed proteomic data from cIV immunocapture.

**Supplemental Table S2.** Processed proteomic data from GFP immunoprecipitation.

**Supplemental Table S3.** Contact probabilities between COA8 and COX10 residues.

**Supplemental Table S4.** Processed proteomic data from complexome profiling analysis.

## METHODS AND PROTOCOLS

### Cell culture conditions

Immortalized human skin fibroblasts and 143B osteosarcoma cells were cultured in DMEM containing 4.5 g/L D-glucose, 110 mg/L sodium pyruvate and GlutaMAX^TM^ supplemented with 10 % FBS. 50 μg/mL uridine was also added to the immortalized fibroblast culture media.

### Lentiviral transduction and immortalization

Primary cultures were immortalized by lentiviral transduction with the pLOX-Ttag-iresTK, obtained from Didier Trono (deposited in Addgene). Constructs expressing WT and mutant human COA8, which were obtained by gene synthesis (GenScript), were cloned into pWPXLd-ires-Puro^R^ lentiviral expression vectors, modified versions of pWPXLd, by restriction enzyme digestion with PmeI and BamHI and ligation using the In-Fusion cloning kit (Takara). Lentiviral particles carrying the construct of interest were packed in HEK293T cells by co-transfection of the pWPXLd vector with the packaging psPAX2 and envelope pMD2.G vectors as described (Perales-Clemente *et al*, 2008). Media containing the lentiviral particles were collected, centrifuged at 5,000 rpm for 5 min at RT and then filtered through 0.45 μm filters. After addition of 5 μg/mL polybrene (hexadimethrine bromide) media were used to treat skin fibroblasts. Twenty-four hours after transduction, cells were selected for puromycin resistance by adding 1 µg/mL puromycin to the cell culture medium.

### In silico protein interaction analyses

Multiple sequence alignments of COA8 and COX10 were generated by using BLAST against a protein database constructed from eukaryotic proteomes (49,102,568 proteins from 2,568 representative or reference genomes from NCBI). To construct the joint alignment for COA8 and COX10, we identified the intersection of the organisms for their orthologous groups and merged the sub-alignments containing only those organisms for the two proteins. Positions that are gaps in the human COA8 and COX10 proteins were removed. AlphaFold was used to model the COA8-COX10 complex based on the joint alignment with a gap of 200 residues introduced in between the alignments. AlphaFold produced probability distribution for residue-residue distances. A contact probability was calculated as the sum of probability for distance bins below 10 Å for each residue pair. The intermolecular residue pairs with a contact probability above 0.9 are considered as interacting residue pairs between COA8 and COX10.

### Blue-Native Gel electrophoresis

Enriched mitochondrial fractions from cells, obtained either by digitonin treatment or by homogenization and differential centrifugation, were solubilized either with 4 mg digitonin per mg of protein or 1.6 mg n-dodecyl-β-D-maltoside (DDM) per mg of protein, and were electrophoresed in precast commercial native gels as described (Fernandez-Vizarra & Zeviani, 2021a).

### Western blot and immunodetection

SDS-PAGE and BN-PAGE gels were transferred to PVDF membranes Tris-Glycine transfer buffer (25 mM Tris-HCl, 192 mM Glycine, 20% methanol, 0.025% SDS) or in Dunn’s carbonate buffer (10 mM NaHCO_3_, 3 mM Na_2_CO_3_), respectively, applying a constant voltage of 100 V at 4°C for 1 hour using a Mini Trans-Blot® Cell (Bio-Rad). For the immunodetection of specific protein targets, blotted PVDF membranes were blocked in 5% skimmed milk in PBS-T (PBS plus 0.1% Tween-20) at RT for 1 hour and then incubated with primary antibodies diluted in 3% BSA in PBS-T O/N at 4°C. The PVDF membranes were washed three times with PBS-T for 10 minutes, incubated with the secondary HRP-conjugated antibody for 1 hour at room temperature and washed three times with PBS-T for 10 minutes before developing. Chemiluminescent signals were recorded using an Alliance Mini HD9 (UVITEC). Alternatively, fluorescent secondary antibodies were used and the signals were visualized using an Odissey Li-Cor system. Antibodies used in this study are listed in the Key Resources Table.

### SILAC and co-immunoprecipitation

The two cell lines to be compared were grown in ‘‘heavy’’ SILAC DMEM (Thermo-Fisher) containing ^15^N– and ^13^C-labeled Arg and Lys and in ‘‘light’’ SILAC DMEM containing ^14^N– and ^12^C– Arg and Lys (Sigma-Aldrich) for at least ten passages. Equal portions of the differentially labeled cell lines were mixed and cells were treated with 4 mg/mL digitonin for 10 minutes on ice to obtain mitochondria-enriched fractions. After two washes with cold PBS, these fractions were then solubilized in native conditions on ice for 10 minutes with PBS containing 10% glycerol, 1.5% DDM, 1X protease inhibitor cocktail and a mixture of polar lipids (Avanti polar lipids), specifically 16:0-18:1 PC (<u>1-palmitoyl-2-oleoyl-glycero-3-phosphocholine)</u>, 16:0-18:1 PE (<u>1-palmitoyl-2-oleoyl-sn-glycero-3-phosphoethanolamine)</u>, 16:0-18:1 PG (<u>1-palmitoyl-2-oleoyl-sn-glycero-3-phospho-(1’-rac-glycerol) (sodium salt))</u>. Samples were centrifuged at 18,000 x *g* for 10 minutes at 4°C. Supernatants were collected and incubated with either monoclonal anti-HA-agarose beads (Sigma), ChromoTek GFP-Trap Agarose beads (Proteintech) or complex IV immunocapture kit (Abcam) O/N at 4°C on a rotating wheel. Beads were washed five times using PBS with 10% glycerol, 0.05% DDM, protease inhibitor cocktail and polar lipids. Immunopurified complexes were eluted under denaturing conditions using 200 mM glycine pH 2.5. After elution samples were neutralized with unbuffered 1 M Tris.

### Mass spectrometry proteomic analysis of immunopurified fractions

After cIV immunocapture, the liquid-free beads were supplemented with 50 µl of reducing loading buffer (12 % SDS, 6 %(v/v) β-mercaptoethanol, 30 %(m/v) glycerol, 0.05 % Coomassie blue G-250, 150 mM Tris-HCl, pH 7.0) and incubated at 45 °C for 15 min to denature and release immunocaptured proteins. Each sample (including beads) was loaded on a 10% Tricine-SDS gel (without stacking layer) and ran for ∼0.5 cm (30 min at 50 V). Precision plus protein standards (BioRad) were used to monitor the correct protein loading into the gel. The gel slices were cut, fixed, destained and processed following in-gel trypsin digestion as detailed in Potter et al (Potter *et al*, 2023). The tryptic peptides were resuspended in 20 µl of 0.5 % formic acid (FA)/5 % acetonitrile (ACN).

The peptides (5 µl of each sample) were injected and analyzed in triplicate using liquid chromatography-tandem mass spectrometry (LC-MS/MS) in a Q Exactive mass spectrometer equipped with an Easy nLC1000 instrument (Thermo Fisher Scientific). The peptides were first loaded on an Acclaim™ PepMap™ 100 C18 LC Column (0.1 mm x 20 mm, nanoViper, 5 µm, 100 Å) and separated using emitter columns (15 cm length × 100 μm ID × 360 μm OD × 15 μm orifice tip; MS Wil/CoAnn Technologies) filled with ReproSilPur C18-AQ reverse-phase beads of 3 μm, 100 Å (Dr. Maisch GmbH). HPLC settings: linear gradients of 5 – 35 % ACN, 0.1 % FA (30 min) at a flow rate of 300 nl·min^-1^, followed by 35 – 80 % ACN, 0.1 % FA (5 min) at 300 nl·min^-1^, and a final column wash with 90 % ACN (5 min) at 600 nl·min^-1^. MS data were collected in Data-dependent acquisition (DDA) mode and positive polarity. The full MS scan range was 300 to 2,000 m/z with a resolution of 70,000 and an automatic gain control (AGC) value of 1E6 with a maximal injection time of 20 ms. The 20 most abundant precursors were selected for MS2. Only charged ions >1 and <5 were selected for MS/MS scans with a resolution of 17,500, an isolation window of 4.0 m/z, and an AGC value set to 1E5 ions with a maximal injection time of 50 ms. (N)CE: 30, dynamic exclusion: 30 s and spectrum data type: profile for both MS1 and MS2.

MaxQuant (v2.0.3.0) (Tyanova *et al*, 2016) was used to analyze the MS raw spectra files. The human reference proteome database (UniProt, May 2022, including canonical sequences and isoforms) was used for identification with a false discovery rate (FDR) ≤1%. N-terminal acetylation, oxidation (M) and deamidation (NQ) were selected as variable modifications. Carbamidomethylation (C) was set as a fixed modification. Specific settings changed: multiplicity: 2; heavy labels: Arg10, Lys8; match between runs: 1 min time window (match unidentified features: true); minimal peptide length: 6; use only unmodified peptides for quantification: false. The MaxQuant list of contaminants was included for the search. All other parameters were kept as default. To quantify protein abundance, label free-quantification (LFQ) and intensity-based absolute quantification (iBAQ) values were calculated. The identified protein groups and peptides were manually inspected and analyzed using Microsoft Excel.

50 µL (15-17 µg) of anti-HA immunoprecipitated protein extract were diluted in 200 µL of washing buffer (WB, Urea 8M, Tris-HCl 100 mM, pH8.5) and loaded into a Vivacon 500 filter (Sartorius, Germany) with a molecular cut-off of 10000 Da to perform a protein digestion based on Filter Aided Sample Preparation (Wisniewski, 2018). Three washes with WB and subsequent centrifugation at 18600 x *g* were performed, followed by the reduction of disulfide bonds with 50 mM dithiothreitol (Fluka) in WB (30 minutes of incubation at 55°C). Samples were centrifuged to discard solutions and the alkylation of cysteines was carried out with 50 mM iodoacetamide (Sigma-Aldrich) in WB (incubation at room temperature for 20 minutes under dark condition). Proteins were finally washed twice with WB and twice with ammonium bicarbonate 100 mM and 50 mM, respectively. Protein digestion was performed overnight at 37°C adding 0.3 µg of sequencing grade modified trypsin (Promega, USA). Three extraction steps performed by washing with 50 mM ammonium bicarbonate and centrifuging at 18600 x *g* for 10 min, allowed to collect peptides that were finally acidified to pH 3 with formic acid, dried under vacuum and stored at −20 °C until mass spectrometry analysis.

Each sample was dissolved in 120 µL of 3% acetonitrile/0.1% formic acid and 8 µL of the resulting solution was injected into a picofrit capillary column (11 cm length, 75 µm internal diameter, New Objectives) packed in house with C18 material (ReproSil, 300 Å, 3 μm; Dr. Maisch HPLC GmbH). Peptides were separated at a flow rate of 250 nL/min using a nanoHPLC Ultimate 3000 (Dionex – Thermo Fisher Scientific) coupled with a LTQ-Orbitrap XL mass spectrometer (Thermo Fisher Scientific), using a linear gradient of acetonitrile/0.1% formic acid from 3% to 40% in 40 min. A Top10 data dependent acquisition method was used for the analysis: a full MS scan (range 300-1700 Da) was performed at high resolution (60000) on the Orbitrap, followed by MS/MS scans on the 10 most intense ions at low resolution in the Linear Ion Trap.

Data were analyzed with the software package Proteome Discoverer 1.4 (Thermo Fisher Scientific) connected to a Mascot search engine (version 2.2.4, Matrix Science) and searched against the Human section of the Uniprot database (version Sept2020, 75074 entries). Trypsin was set as proteolytic enzyme with up to two missed cleavages allowed. Carbamidomethylation was set as fixed modification while methionine oxidation, ^13^C_6_^15^N_2_ Lysine and ^13^C_6_^15^N_4_ Arginine were set as variable modifications. The Percolator algorithm was used to assess the False Discovery Rate (FDR), proteins were grouped into families according to the principle of maximum parsimony, and results were filtered to keep into account only proteins identified with at least 2 unique peptides with high confidence (FDR < 0.01). The heavy to light ratios (H/L) were computed by the software as the median value of the H/L of all peptides belonging to the same protein.

### Enzyme activity assays

Cytochrome c oxidase (COX) and citrate synthase (CS) activities were measured in digitonin-treated cells by spectrophotometric kinetic assays in a 96-well plate reader, as described (Brischigliaro *et al*, 2022; Tiranti *et al*, 1995).

### Isolation of mitochondria

Mitochondria from patient-derived immortalized skin fibroblasts were released by hypotonic shock and isolated using differential centrifugation. Cellular pellets were diluted in PBS and centrifuged at 600 x *g* (10 min, 4° C). Supernatant was removed, pellet was resuspended 10:1 (v/w) in a hypotonic solution of 10 mM Tris pH 7.6, supplemented with protease inhibitor cocktail (PIC). Immediately, suspension was homogenized with Dounce homogenizer (20 strokes, loose pestle), sucrose was added to the homogenate (1.5 M stock solution, final concentration = 250 mM, supplemented with PIC) and then followed a centrifugation at 600 x *g* (10 min, 4°C). Supernatant was recovered into a clean tube and centrifuged at 10,000 x *g* (10 min, 4°C). Pelleted mitochondria were resuspended in 500 µL of SEKT (250 mM sucrose, 40 mM KCl, 20 mM Tris-HCl, 2mM EGTA, pH 7.,6) with PIC and centrifuged again at 10,000 x *g* (10 min, 4°C). The final mitochondrial fractions were stored as dry pellets at –80°C.

### Quantification of mitochondrial hemes

200 µL of acetone containing 2.5% HCl were added to 500 µg of mitochondrial protein, vortexed, and centrifuged (10 min, 15,000 x *g*, 4°C). After, 200 µL of water/acetonitrile 50/50 were added to the supernatant, and pH was adjusted to 3.5 by adding approximately 5 µL of 30% ammonium hydroxide. After centrifugation (10 min, 15,000 x *g*, 4°C), 100 µL of the supernatant were analyzed by UHPLC-HRMS.

The UHPLC-HRMS system was equipped with Agilent 1260 Infinity II LC System coupled to a DAD and an Agilent 6545 Q-TOF mass analyzer (Agilent Technologies, Palo Alto, Santa Clara, CA, USA). The analytical column was a Kinetex 2.6 μm C18 Polar, 100 A, 100 × 2.1 mm (Phenomenex, Bologna, Italy), thermostated at 45 °C. The components of the mobile phase A and B were water and acetonitrile, respectively, both containing 0.1% trifluoroacetic acid. The eluent flow rate was 0.4 mL/min. The mobile phase gradient profile was as follows (t in min): t0–10 30%-100% B; t10–15 100% B, t15–22 30% B. DAD signals were recorded in the 210-600 nm range, and chromatograms were extracted at (400 ± 4) nm.

The MS conditions were: electrospray (ESI) ionization in positive mode, gas temperature 325 °C, drying gas 11 L/min, nebulizer 45 psi, sheath gas temperature 275 °C, sheath gas flow 12 L/min, VCap 4000 V, nozzle voltage 0 V, fragmentor 180 V. Centroid full scan mass spectra were recorded in the range 100–1700 m/z with a scan rate of 1 spectrum/s. MS/MS data were acquired in targeted mode with a scan rate of 2 spectrum/s, collision energy of 30 eV, and isolation width of 1.3 Da. The QTOF calibration was performed daily with the manufacturer’s solution in this mass range. The MS and MS/MS data were analyzed by the Mass Hunter Qualitative Analysis software (Agilent Technologies, Palo Alto, Santa Clara, CA, USA). The MS chromatographic peaks of heme A and heme B were identified and integrated by considering the extracted ion chromatogram (EIC) of the [M + H]^+^ species (*m/z* 852.3565 for heme A and *m/z* 616.1768 for heme B, respectively) selected with a window of 5 ppm. Identification of heme A and heme B were also confirmed by MS/MS data.

### Complexome profiling

After BN-PAGE, the gel was stained with Coomassie Blue and further scanned to prepare a template for cutting. Each lane was cut into 48 fractions and collected in 96 well-filter plates. The gel pieces were washed several times in 60% methanol, 50 mM ammonium bicarbonate (ABC). Excess solution was removed by centrifugation for 2 min at 600 X g. Proteins were reduced in 10 mM DTT, 50 mM ABC for 1 h at 56 °C and alkylated for 45 min in 30 mM iodoacetamide, 50 mM ABC. Proteins were digested for 16 hours with trypsin (sequencing grade, Promega) at 37 °C in 50 mM ABC, 0.01 % Protease Max (Promega), 1 mM CaCl_2_. Tryptic peptides were eluted with 30 % ACN/3 % FA and recovered by centrifugation in a 96 wells microplate, speed vac-dried and resuspended in 15 µl of 1% ACN and 0.5% FA. Dissolved peptides were stored at –20 °C until MS analysis.

LC-MS/MS analysis was performed using a Q Exactive Plus mass spectrometer equipped with an UHPLC Dionex Ultimate 3000 instrument (Thermo Fisher Scientific). The peptides (3 µl of each fraction) were loaded on an Acclaim™ PepMap™ 100 C18 LC Column (0.1 mm x 20 mm, nanoViper, 5 µm, 100 Å) and separated using emitter columns (15 cm length × 100 μm ID × 360 μm OD × 15 μm orifice tip; MS Wil/CoAnn Technologies) filled with ReproSilPur C18-AQ reverse-phase beads of 3 μm, 100 Å (Dr. Maisch GmbH). Each fraction was analyzed twice. HPLC settings: linear gradients of 4 – 25 % ACN, 0.1 % FA for 35 min followed by 25 – 50 % ACN, 0.1 % FA for 5 min, 50 – 99 % ACN, 0.1% FA for 1 min. The column was washed with 99 % ACN, 0.1% FA for 5 min and then equilibrated with 4 % ACN, 0.1 % FA for 14 min. All flow rates were set as 300 nl·min^-1^. MS data were recorded by DDA. The full MS scan range was 300 to 2,000 m/z with a resolution of 70,000 and an AGC of 3E6 with a maximal injection time of 65 ms. The 20 most abundant precursors were selected for MS2. Only charged ions >2 and <8 were selected for MS/MS scans with a resolution of 17,500; isolation window of 2.0 m/z; AGC: 1E5; maximal injection time: 65 ms. MS data were acquired in profile mode.

MaxQuant 2.4.13.0 (Tyanova *et al*., 2016) was used to analyze the MS raw spectra files. The human reference proteome database (UniProt, February 2024, including canonical sequences and isoforms) was used for identification with a false discovery rate (FDR) ≤1%. N-terminal acetylation, oxidation (M) and deamidation (NQ) were selected as variable modifications. Carbamidomethylation (C) was set as a fixed modification. Specific settings changed: match between runs: 1 min time window (match unidentified features: true); minimal peptide length: 7; max. missed cleavages: 2; use only unmodified peptides for quantification: false. The MaxQuant list of contaminants was included for the search. All other parameters were kept as default. To quantify the protein abundances, iBAQ values were calculated. The proteinGroups.txt output file was automatically processed using “process_maxquant” (https://github.com/joerivstrien/process_maxquant). The resulting dataset was manually inspected and analyzed using Microsoft Excel. The list of protein groups was hierarchically clustered based on the abundance patterns with an average linkage algorithm (absolute correlation) using Cluster 3.0 (de Hoon *et al*, 2004). Individual profiles were normalized using the sum of iBAQ values of the identified proteins annotated in MitoCarta 3.0 (Rath *et al*, 2021). Calibration curves of apparent molecular masses were generated using human membrane and water-soluble protein complexes of known masses. The normalized profiles derived from the replicates of the same condition were averaged. Averaged profiles were normalized to the maximal iBAQ value across all slices for each protein group to generate the relative abundance profiles. The data were visualized as heatmaps and line charts generated in Microsoft Excel.

## QUANTIFICATION AND STATISTICAL ANALYSIS

Biochemical experiments were performed in triplicate and the graphical results are presented as mean ± standard deviation (SD). The full data distribution in the graphs is shown overlaying the dot plots, corresponding to the individual values, to the bar graphs. To assess the statistical significance of the mean differences, one-way ANOVA tests with either Tukey’s or Dunett’s post-hoc tests for multiple comparisons were applied, following the recommendations of the GraphPad Prism software (versions 8 and 10). P values < 0.05 were considered significant.

KEY RESOURCES TABLE.

KEY RESOURCES TABLE
| REAGENT or RESOURCE | SOURCE | IDENTIFIER |
| --- | --- | --- |
| **Antibodies** |  |  |
| Anti-SDHB antibody | Abcam | Cat#ab14714, RID:AB_301432 |
| Anti-GFP antibody | Abcam | Cat#ab1218, RRID:AB_298911 |
| Anti-COX10 antibody | Atlas Antibodies | Cat#HPA032005, RRID:AB_10603560 |
| Anti-Cytochrome C oxidase subunit II antibody | Abcam | Cat#ab110258, RRID:AB_10887758 |
| Anti-SDHA Monoclonal Antibody | Abcam | Cat#ab14715, RRID:AB_301433 |
| Anti-β-Actin Antibody | Santa Cruz Biotechnology | Cat#sc-47778, RRID:AB_626632 |
| Anti-NDUFB8 antibody | Proteintech | Cat#14794-1-AP; RRID:AB_2150970 |
| Anti-UQCRC1 antibody | Proteintech | Cat#21705-1-AP, RRID:AB_10734437 |
| Anti-UQCRC2 antibody | Proteintech | Cat#14742-1-AP, RRID:AB_2241442 |
| Anti-ATP5A1 antibody | Proteintech | Cat#14676-1-AP, RRID:AB_2061761 |
| Anti-MTCO1 antibody | Abcam | Cat#ab14705, RRID:AB_2084810 |
| Monoclonal Anti-HA-Agarose | Merck | Cat#A2095 |
| ChromoTek GFP-Trap® Agarose | Proteintech | Cat#gta |
| Complex IV Immunocapture Kit | Abcam | Cat#ab109801 |
| Goat anti-Rabbit IgG (H+L) Highly Cross-Adsorbed Secondary Antibody, Alexa Fluor™ Plus 800 | Thermo Fisher Scientific | Cat#A32735, RRID AB_2633284 |
| Goat anti-Mouse IgG (H+L) Cross-Adsorbed Secondary Antibody, Alexa Fluor™ 680 | Thermo Fisher Scientific | Cat#A-21057, RRID AB_2535723 |
| Anti-Mouse IgG (H+L), HRP Conjugate | Promega | Cat#W4021, RRID:AB_430834 |
| Anti-Rabbit IgG (H+L), HRP Conjugate | Promega | Cat#W4011, RRID:AB_430833 |
| **Bacterial and virus strains** |  |  |
| **Biological samples** |  |  |
| **Chemicals, peptides, and recombinant proteins** |  |  |
| 16:0-18:1 PC | Avanti Polar Lipids | Cat#850457C |
| 16:0-18:1 PE | Avanti Polar Lipids | Cat#850757C |
| 16:0-18:1 PG | Avanti Polar Lipids | Cat#840457C |
| Acetyl coenzyme A lithium salt | Merck | Cat#A2181 |
| 6-Aminocaproic acid | Merck | Cat#A2504 |
| BSA (fatty acid free) | Merck | Cat#A6003 |
| Cytochrome c from equine heart | Merck | Cat#C7752 |
| cOmplete™, EDTA-free Protease Inhibitor Cocktail | Roche | Cat#04693132001 |
| DMEM for SILAC | Thermo Fisher Scientific | Cat#88364 |
| DMEM, high glucose, GlutaMAX™ Supplement, pyruvate | Thermo Fisher Scientific | Cat#31966047 |
| DPBS, no calcium, no magnesium | Thermo Fisher Scientific | Cat#14190169 |
| DTNB (5,5'-Dithiobis(2-nitrobenzoic acid)) | Merck | Cat#D218200 |
| n-Dodecyl-beta-maltoside (DDM) | Merck | Cat#D4641 |
| Digitonin, High Purity | Calbiochem | Cat#300410 |
| EDTA (Ethylenediaminetetraacetic acid disodium salt dihydrate) | Merck | Cat#E1644 |
| EGTA (Ethylene glycol-bis(2-aminoethylether)-N,N,N',N'-tetraacetic acid) | Merck | Cat#E3889 |
| Enhanced Chemiluminescent Substrate | EuroClone | Cat#EMP001005 |
| Fetal Bovine Serum, qualified, Brazil | Thermo Fisher Scientific | Cat#10270106 |
| Glycine | Merck | Cat#G8898 |
| HBSS, no calcium, no magnesium | Thermo Fisher Scientific | Cat#14170088 |
| L-Lysine-<sup>13</sup>C<sub>6</sub>, <sup>15</sup>N<sub>2</sub> hydrochloride | Merck | Cat#608041 |
| L-Arginine-<sup>13</sup>C<sub>6</sub>, <sup>15</sup>N<sub>4</sub> hydrochloride | Merck | Cat#608033 |
| Methanol | Merck | Cat#34860-2.5L-R |
| MOPS | Merck | Cat#M1254 |
| NativePAGE Cathode Buffer Additive (20X) | Thermo Fisher Scientific | Cat#BN2002 |
| NativePAGE Running Buffer (20X) | Thermo Fisher Scientific | Cat#BN2001 |
| NuPAGE LDS Sample Buffer (4X) | Thermo Fisher Scientific | Cat#NP0007 |
| NuPAGE MOPS SDS Running Buffer (20X) | Thermo Fisher Scientific | Cat#NP0001 |
| Oxaloacetic acid | Merck | Cat#O4126 |
| PBS Tablets | Merck | Cat#524650-1EA |
| Potassium chloride | Merck | Cat#P3911 |
| Potassium phosphate dibasic | Merck | Cat#P2222 |
| Potassium phosphate monobasic | Merck | Cat#P5655 |
| Prestained Protein SHARPMASSTM VI | EuroClone | Cat#EPS025500 |
| Puromycin | InvivoGen | Cat#ant-pr-1 |
| SERVA Blue G | Serva | Cat#35050.02 |
| Sodium carbonate | Merck | Cat#S7795 |
| Sodium bicarbonate | Merck | Cat#S8875 |
| Sodium dodecyl sulphate | Merck | Cat#L3771 |
| Sucrose | Merck | Cat#S7903 |
| TRIS base | Merck | Cat#T1503 |
| Triton X-100 | Merck | Cat#T8787 |
| Trypsin-EDTA (0.5%), no phenol red | Thermo Fisher Scientific | Cat#15400054 |
| Tween-20 | Merck | Cat#P7949 |
| Uridine | Merck | Cat#U3003 |
| **Critical commercial assays** |  |  |
| **Deposited data** |  |  |
| Complexome Profiling data | CEDAR (van Strien *et al.*, 2021) (https://www3.cmbi.umcn.nl/cedar/browse/) | XXXX |
| **Experimental models: Cell lines** |  |  |
| S2 primary fibroblasts | Daniele Ghezzi (Fondazione IRCCS Istituto Neurologico Carlo Besta, Milan, Italy) | Melchionda *et al.* (2014)(Melchionda *et al.*, 2014) |
| S6 primary fibroblasts | Rosalba Carrozzo (Bambino Gesù Children's Hospital, IRCCS, Rome, Italy) | Melchionda *et al.* (2014)(Melchionda *et al.*, 2014) |
| S2 immortalized fibroblasts | Erika Fernandez-Vizarra | Signes *et al.* (2019)(Signes *et al.*, 2019) |
| S6 immortalized fibroblasts | Erika Fernandez-Vizarra | Signes *et al.* (2019)(Signes *et al.*, 2019) |
| S6+GFP | Erika Fernandez-Vizarra | Signes *et al.* (2019)(Signes *et al.*, 2019) |
| S6+COA8<sup>GFP</sup> | Erika Fernandez-Vizarra | Signes *et al.* (2019)(Signes *et al.*, 2019) |
| S6+F118A<sup>GFP</sup>, F118S<sup>GFP</sup>, F159A<sup>GFP</sup>, W174A<sup>GFP</sup> | Erika Fernandez-Vizarra | This work |
| 143B+GFP | Erika Fernandez-Vizarra | Signes *et al.* (2019)(Signes *et al.*, 2019) |
| 143B+COA8<sup>GFP</sup> | Erika Fernandez-Vizarra | Signes *et al.* (2019)(Signes *et al.*, 2019) |
| 143B+COA8<sup>HA</sup> | Erika Fernandez-Vizarra | Signes *et al.* (2019)(Signes *et al.*, 2019) |
| **Experimental models: Organisms/strains** |  |  |
| **Oligonucleotides** |  |  |
| **Recombinant DNA** |  |  |
| pLOX-Ttag-iresTK | Addgene | Addgene #12246 |
| pWPXLd | Addgene | Addgene #12258 |
| psPAX2 | Addgene | Addgene #12260 |
| pMD2.G | Addgene | Addgene #12259 |
| COA8<sup>HA</sup>-pWPXLd-ires-Puro<sup>R</sup> | Erika Fernandez-Vizarra | Signes et al. (2019)(Signes *et al.*, 2019) |
| GFP-pWPXLd-ires-Puro<sup>R</sup> | Erika Fernandez-Vizarra | Signes et al. (2019)(Signes *et al.*, 2019) |
| COA8<sup>GFP</sup>-pWPXLd-ires-Puro<sup>R</sup> | Erika Fernandez-Vizarra | Signes et al. (2019)(Signes *et al.*, 2019) |
| F118A<sup>GFP</sup>-pWPXLd-ires-Puro<sup>R</sup> | Erika Fernandez-Vizarra | This work |
| F118S<sup>GFP</sup>-pWPXLd-ires-Puro<sup>R</sup> | Erika Fernandez-Vizarra | This work |
| F159A<sup>GFP</sup>-pWPXLd-ires-Puro<sup>R</sup> | Erika Fernandez-Vizarra | This work |
| W174A<sup>GFP</sup>-pWPXLd-ires-Puro<sup>R</sup> | Erika Fernandez-Vizarra | This work |
| **Software and algorithms** |  |  |
| Adobe Photoshop 2024 | Adobe | RRID:SCR_014199 |
| CorelDRAW 2017 | Corel Corporation | RRID:SCR_013674 |
| Graphpad Prism v.8 and 10 | GraphPad Software | RRID:SCR_000306 |
| Image Lab Software | BioRad | RRID:SCR_014210 |
| Jalview | http://www.jalview.org/ | RRID:SCR_006459 |
| Proteome Discoverer v.1.4 | Thermo Fisher Scientific | RRID:SCR_014477 |
| MaxQuant v.2.0.3.0 and 2.4.13.0 | https://www.maxquant.org/ | RRID:SCR_014485 |
| Maxquant output processing for Complexome Profiling | Ariel Komen and Joeri van Strien | https://github.com/joerivstrien/process_maxquant |
| **Other** |  |  |
| NativePAGE 3-12% Bis-Tris Gels | Thermo Fisher Scientific | Cat#BN1001BOX |
| NativePAGE 4-16% Bis-Tris Gels | Thermo Fisher Scientific | Cat#BN1002BOX |
| NuPAGE™ 12%, Bis-Tris, 1.0 mm | Thermo Fisher Scientific | Cat#NP0341BOX |
| NuPAGE™ 4 to 12%, Bis-Tris, 1.0–1.5 mm | Thermo Fisher Scientific | Cat#NP0326BOX |
| PVDF Transfer Membrane, 0.45 µm, 26.5 cm x 3.75 m | Thermo Fisher Scientific | Cat#88518 |
| XCell SureLock™ Mini-Cell | Thermo Fisher Scientific | Cat#EI0001 |

